# Uncovering the spatiotemporal structure of neural avalanches through optimal transport and dynamic time warping

**DOI:** 10.64898/2026.07.10.737743

**Authors:** Filip Novicky, Nikola Jajcay, Jaroslav Hlinka

**Affiliations:** Department of Complex Systems, Institute of Computer Science of the Czech Academy of Sciences

**Keywords:** Neural Avalanches, Spatiotemporal Patterns, Optimal Transport, Dynamic Time Warping, Psilocybin, EEG, Travelling Waves, Microstates

## Abstract

Neural avalanches—or threshold-defined bursts of coordinated activity—are traditionally characterised by scale-free statistics. However the study of their detailed spatiotemporal structure and their recurrence within spontaneous activity has been hindered by the variability of avalanches in duration and spatial extent. To tackle this challenge, we propose the use of flexible alignment: we employ a distance metric combining unbalanced optimal transport with subsequence dynamic time warping, enabling comparison across events of different lengths and spatial configurations. Applied to 64-channel EEG from 63 participants in the PsiConnect psilocybin study, hierarchical clustering revealed 12 recurring propagation patterns. These cluster templates were then traced in the original continuous recordings, identifying sequences where the same pattern recurred consecutively and immediately at least twice. Sequences with alternating polarity were classified as oscillating; those with consistent polarity as stable. Oscillating sequences predominantly corresponded to clusters exhibiting visually confirmed spatial propagation, while stable sequences corresponded to spatially fixed patterns. Under psilocybin, oscillating sequences were reduced relative to stable sequences, shifting the polarity balance toward stable; this overall shift was confirmed by a subject-level permutation test, while the apparent task- and training-specific effects did not survive it. This effect was also mostly driven by a specific cluster, suggesting that there is concrete neural dynamics that are temporally affected by the consumption of psilocybin. The developed methodology has been implemented in a publicly available stppy Python package.

**Author Summary:** When large groups of neurons activate together in brief bursts—called neural avalanches—they create patterns that sweep across the brain’s surface. These patterns are usually studied only as statistics (how big, how frequent), giving little attention to what they actually look like. We developed a method to compare the shapes of individual avalanches, group similar ones together, and then use the resulting templates as detectors to track when each pattern reappears in continuous brain recordings. Applied to high-density EEG data from 63 human subjects, we discovered 12 propagation patterns, and analysed whether they oscillate (i.e., repeat with alternating polarity) or remain stable, maintaining a fixed spatial arrangement. We found that during psilocybin intoxication, oscillating patterns were suppressed relative to stable patterns, making stable patterns relatively more prevalent in the detected sequences. This overall shift was statistically reliable across participants, although the apparent differences between individual tasks were not. Our approach bridges three previously separate research areas—avalanche dynamics, EEG microstates, and travelling waves—suggesting they describe the same underlying brain activity from different angles: some of our clusters resemble canonical microstate topographies, while others reveal the travelling wave dynamics that microstate analysis, by design, cannot sufficiently capture.

## 1 Introduction

Neural recordings are long, and resting-state EEG in particular offers no external temporal structure to guide analysis (e.g., no stimulus onset, no response window, no task boundary). Most approaches handle this by decomposing the full recording into global patterns (power spectra, functional connectivity, microstate sequences) that collectively explain as much variance as possible. But if coordinated neural activity occurs in brief, isolated episodes rather than continuously, global decompositions will blur these episodes with intervening unstructured activity, obscuring the spatiotemporal organization of the events themselves. A different strategy is to begin from discrete events: extract them, examine their individual spatiotemporal structure, and ask whether similar patterns recur and following what rules—in other words mapping the repertoire and dynamics of brain states.

Neural avalanches—threshold-defined bursts of coordinated activity that begin when signals exceed a statistical threshold and end when activity returns to baseline (Beggs and Plenz, 2003; Plenz et al., 2021)—provide natural candidates for such events. They can be extracted systematically from recordings without imposing external structure, and each avalanche carries its own spatiotemporal shape: a specific spatial configuration of active electrodes evolving over a specific duration. Yet avalanche research has overwhelmingly focused on scale-free statistics and criticality (Friedman et al., 2012; Shew et al., 2009), treating avalanches as points in a size-duration distribution rather than as individual objects with identifiable propagation structure. Whether recurring spatiotemporal motifs exist among avalanches—and if so, what they look like—has not been systematically addressed, largely because comparing avalanches of different durations and spatial extents is methodologically challenging.

We address this by combining Unbalanced Optimal Transport (UOT) with subsequence Dynamic Time Warping (subDTW) to define a pairwise distance between avalanches that respects both spatial structure and temporal dynamics while accommodating minor differences in length and coverage. This approach follows the philosophy of Janati et al. (2020), who combined unbalanced optimal transport with soft-DTW for spatiotemporal alignment. We adapt their framework to the specific geometry of EEG avalanches: the Hungarian algorithm replaces the Sinkhorn solver for simplicity given discrete activations, polarity constraints preserve the physiologically meaningful sign structure of EEG deflections, and subsequence DTW replaces soft-DTW to handle the common case where a shorter avalanche is embedded within a longer one. The resulting distance matrix enables hierarchical clustering into a suitable finite set of key propagation modes, whose consensus templates can then be tracked in continuous recordings through spatiotemporal correlation.

We apply this framework to 64-channel EEG data from 63 participants in the PsiConnect psilocybin study (Novelli et al., 2026), recorded during resting state, meditation, music listening, and video viewing under both baseline and acute psilocybin conditions. Psychedelic states are commonly characterised by increased entropy in neural recordings (Carhart-Harris et al., 2014; Siegel et al., 2024), a global statistic indicating reduced predictability but not specifying what changed structurally. By tracking specific spatiotemporal patterns, we can ask a sharper question: which recurring propagation modes does psilocybin suppress, preserve, or alter?

This work offers three contributions. First, a distance metric and analysis pipeline for comparing avalanches of arbitrary duration and spatial extent, enabling clustering and template-based tracking. Second, we provide a detailed description of individual avalanches that possess identifiable spatiotemporal propagation templates with systematic spatial symmetries. Third, a demonstration that psilocybin shifts the polarity balance of recurring sequences away from oscillating patterns toward stable, spatially fixed patterns—an overall effect whose direction is consistent across resting state, meditation, and music listening. The methodology is available as the stppy Python package at https://github.com/filipnovicky/stppy/.

## 2 Results

### 2.1 Spatiotemporal anatomy of individual clusters

From 5108 avalanches representing 400.44 seconds of data (0.3% of the entire dataset), where each was then supplemented by its reversed-polarity-copy to yield 10216 avalanches for analysis, we were able to identify the following 12 propagation templates, representing clusters with at least 30 avalanches, with an arbitrary distance threshold of 3.5 (see Methods). The templates corresponding to these clusters are shown in Figure 1. Clusters are labeled 0–11 throughout the manuscript for consistency with the analysis pipeline. Of the 5108 original avalanches, 579 were assigned to these selected clusters. Summary statistics for the clusters are provided in Table 1.

**Figure 1:**
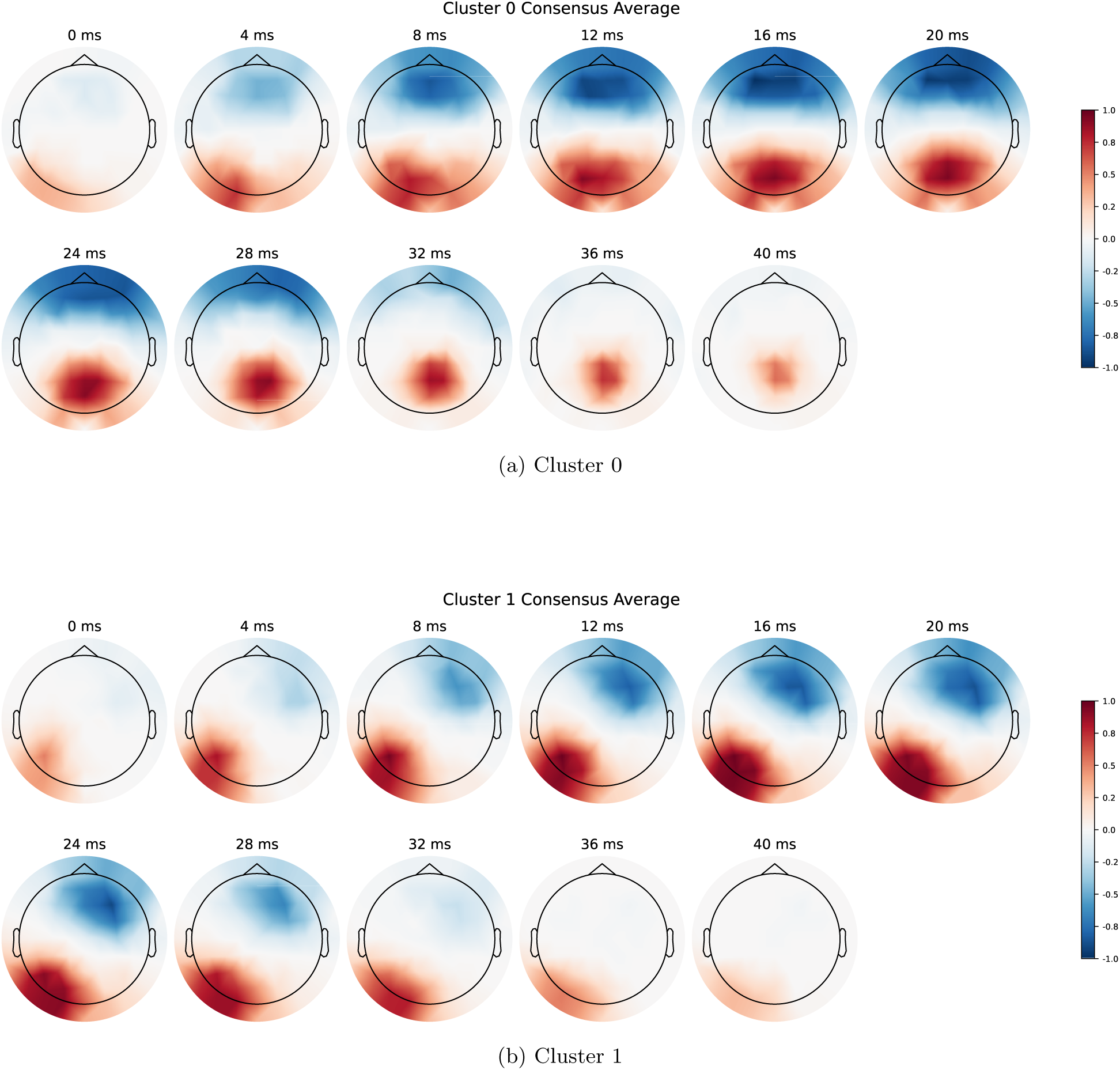
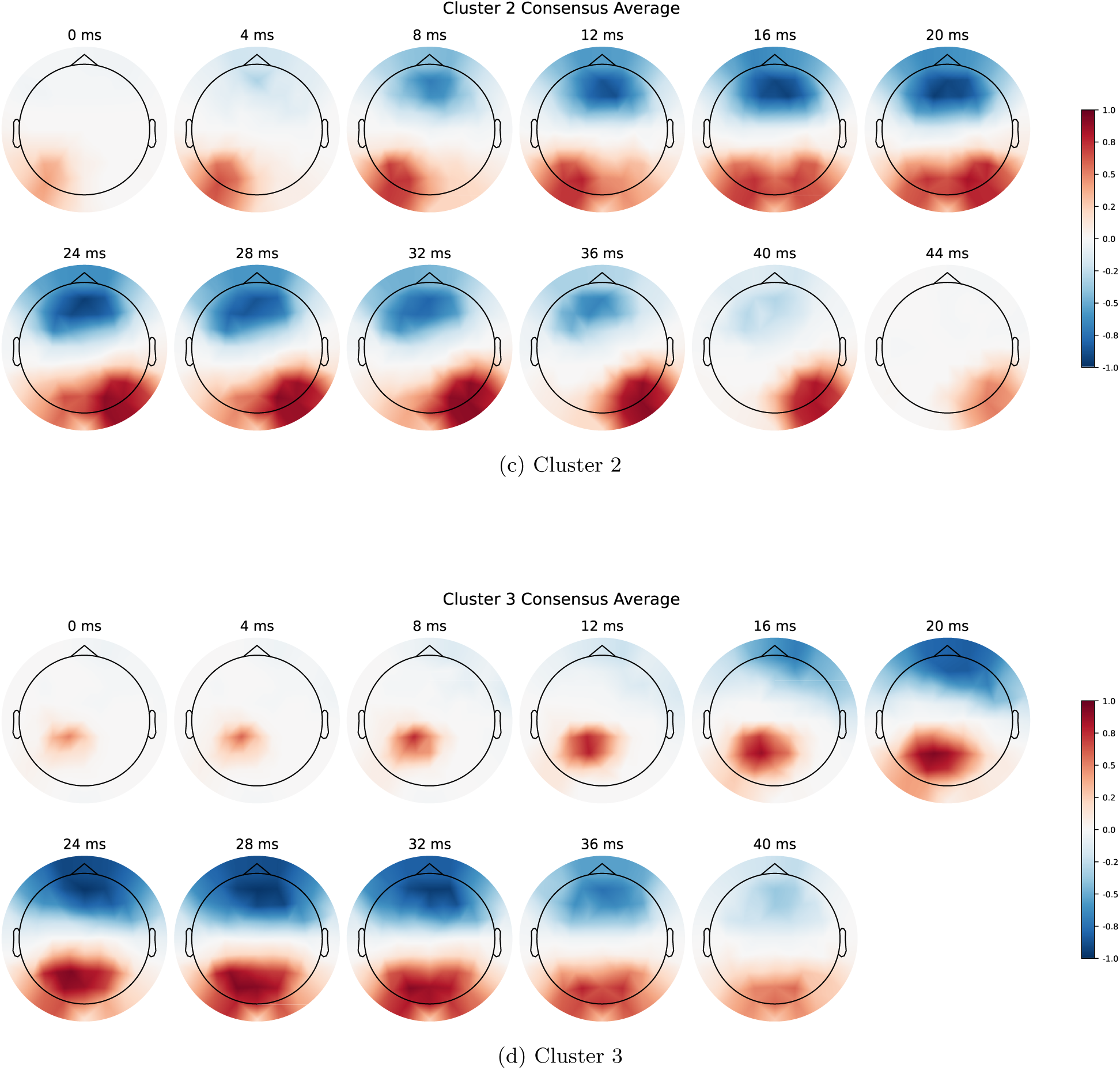
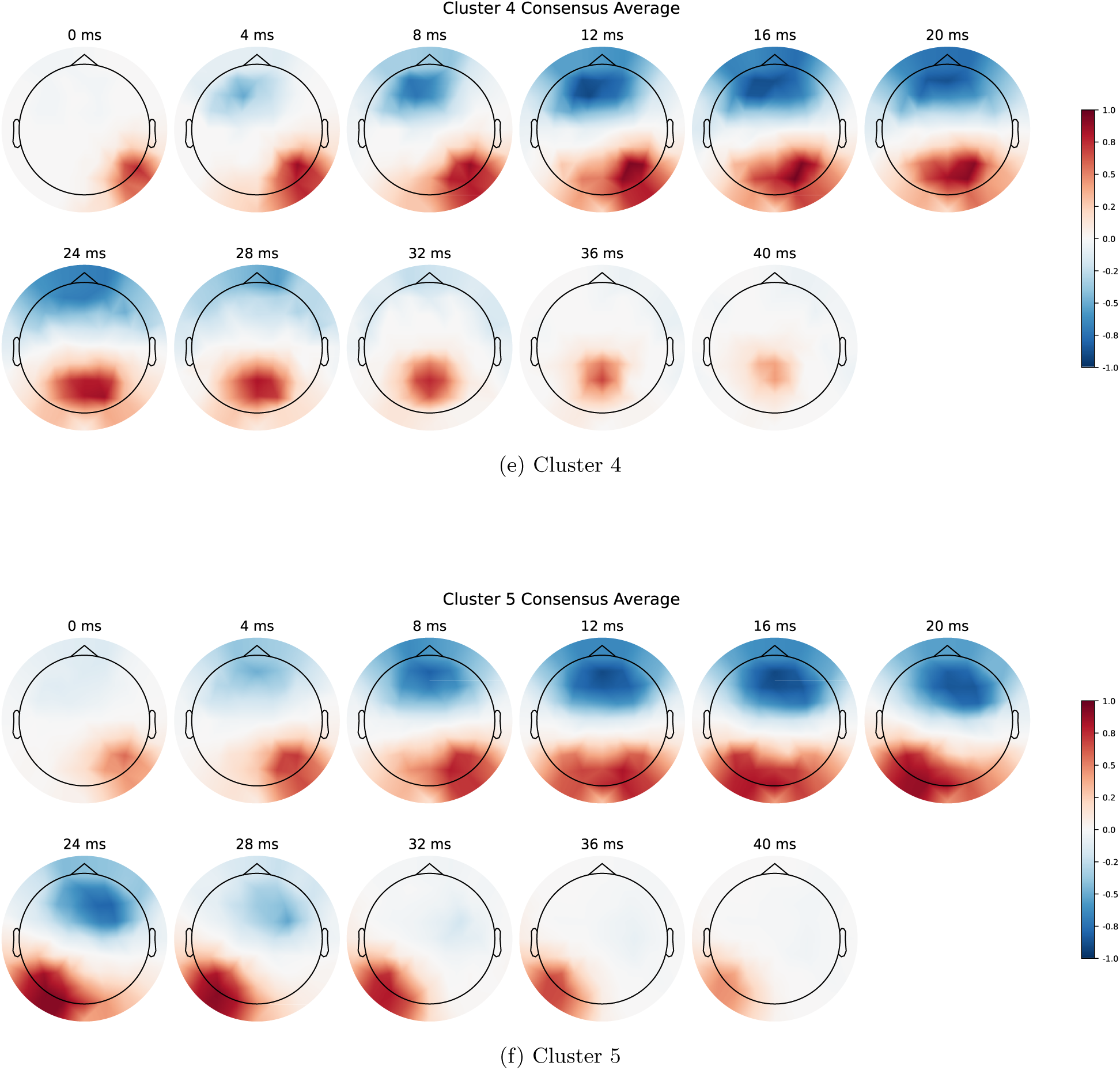
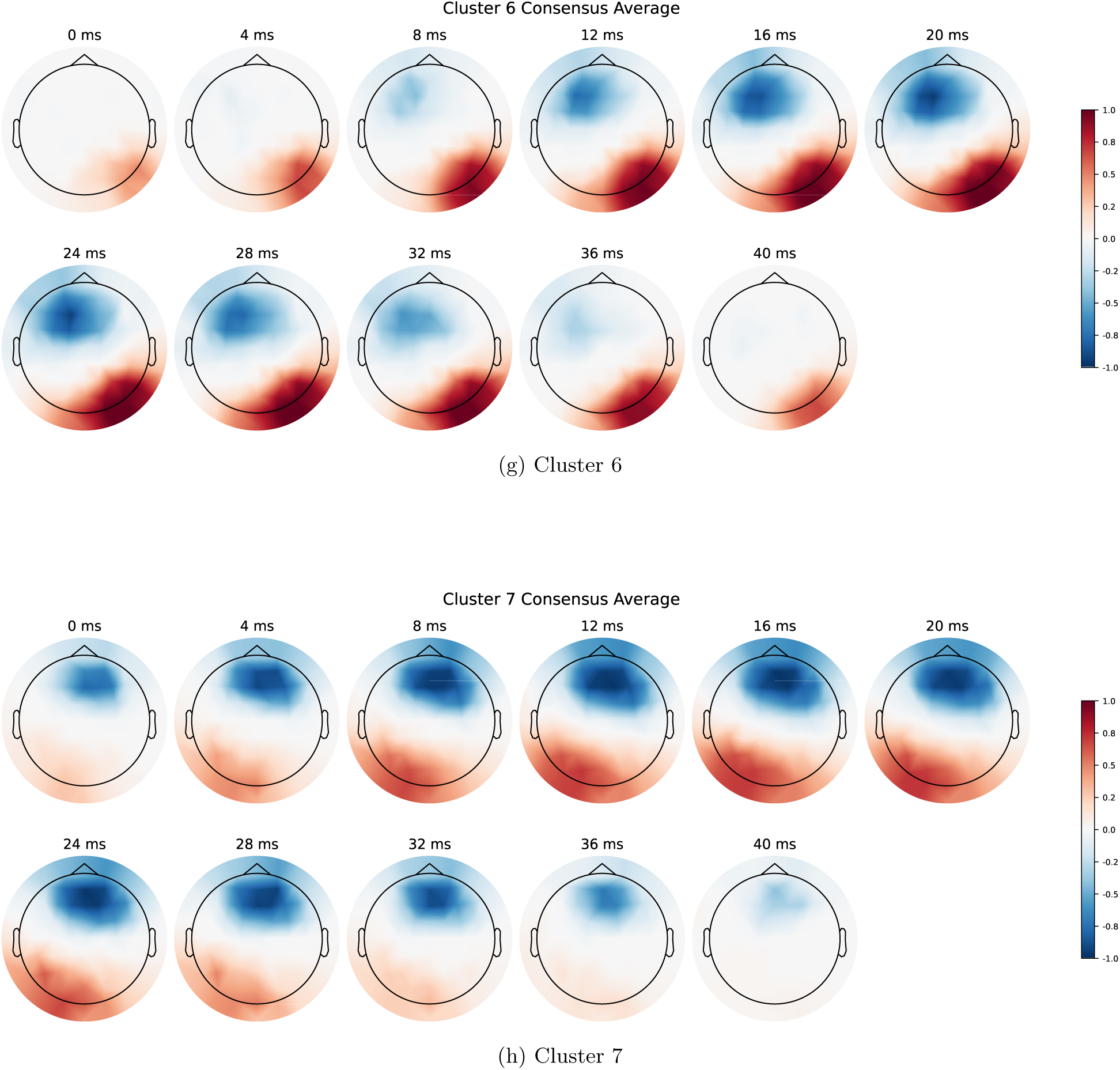
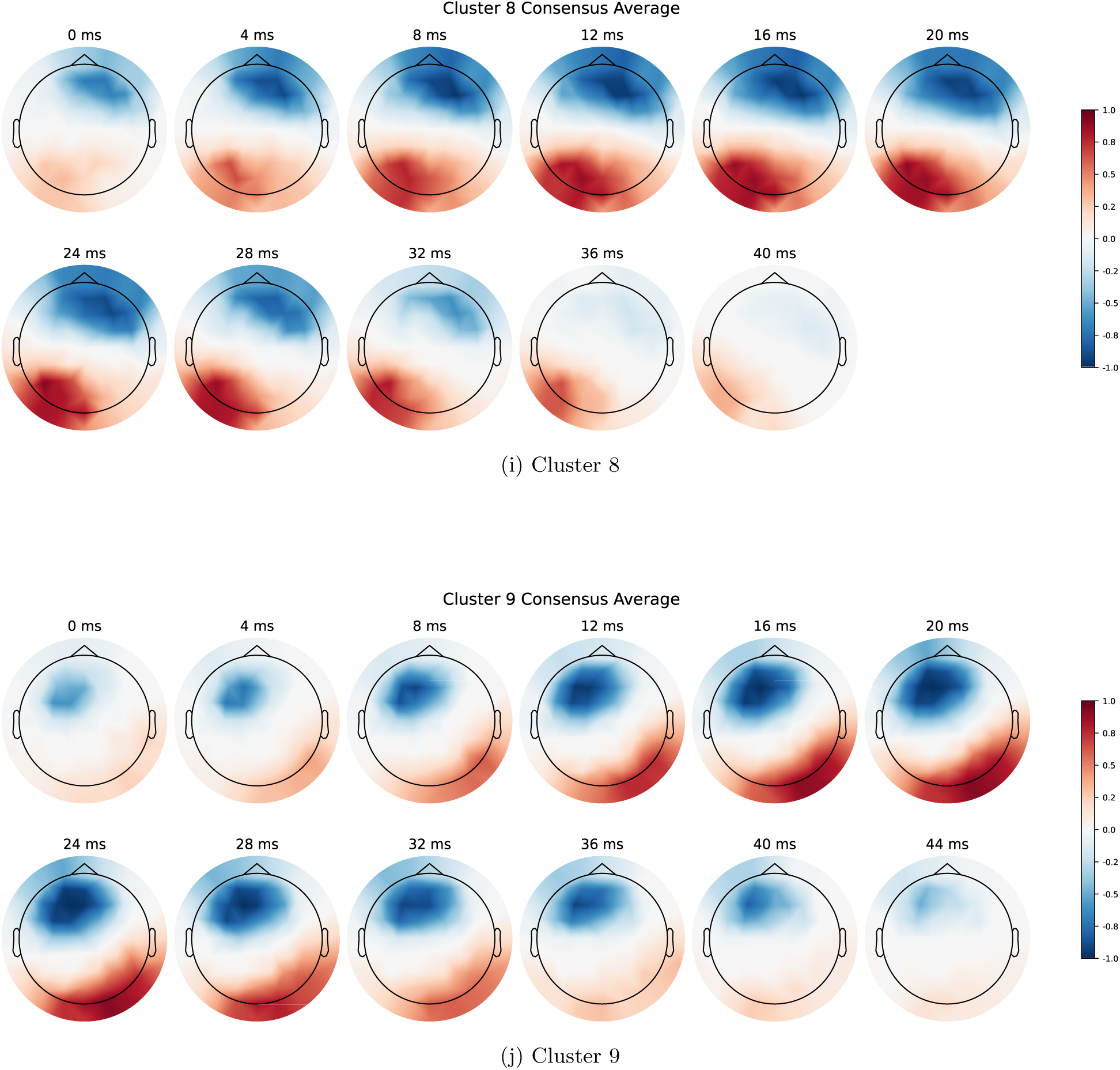
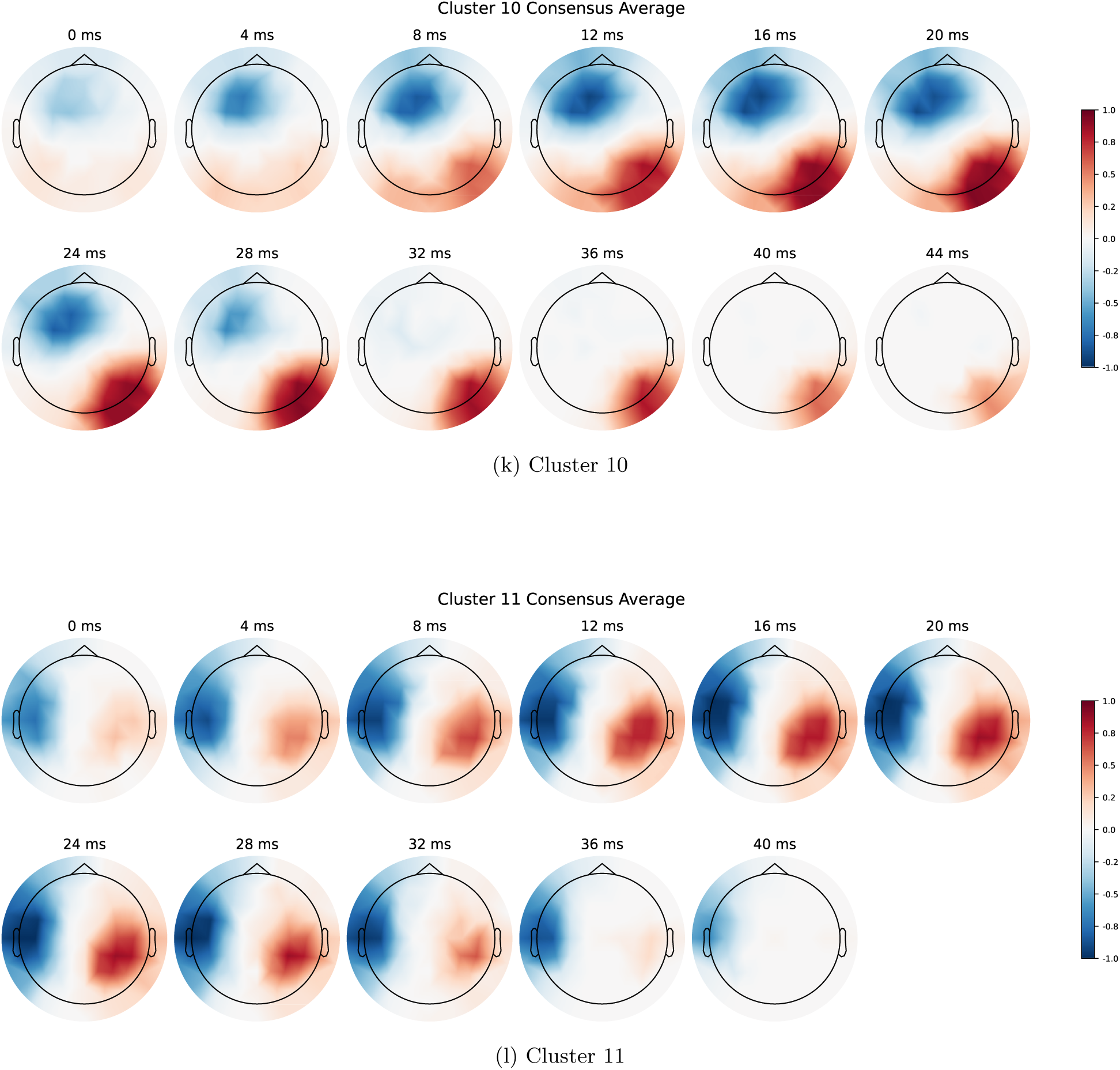
Spatiotemporal avalanche clusters. Each row shows the consensus pattern for one cluster, displaying the temporal evolution of activity across the scalp from left to right. Red indicates positive deflections, blue indicates negative deflections. Time labels above each topomap indicate milliseconds from avalanche onset.

**Table 1:**
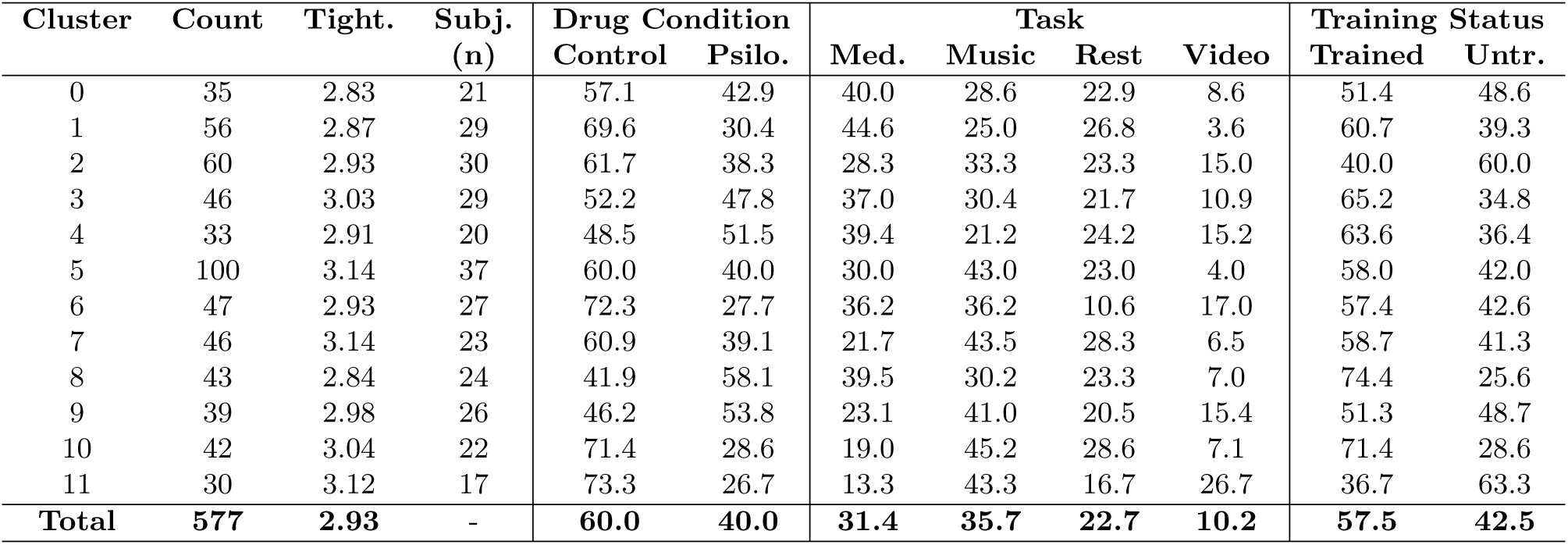
Cluster composition showing distribution of drug conditions, tasks, and training status (values in %). The subject column represents in how many participants the avalanches were observed. The statistics showed that there was no statistically significant difference in drug condition, task or training status after the Benjamini-Hochberg FDR correction.

| Cluster | Count | Tight. | Subj.<br>(n) | Drug Condition |  | Task |  |  |  | Training Status |  |
| --- | --- | --- | --- | --- | --- | --- | --- | --- | --- | --- | --- |
|  |  |  |  | Control | Psilo. | Med. | Music | Rest | Video | Trained | Untr. |
| 0 | 35 | 2.83 | 21 | 57.1 | 42.9 | 40.0 | 28.6 | 22.9 | 8.6 | 51.4 | 48.6 |
| 1 | 56 | 2.87 | 29 | 69.6 | 30.4 | 44.6 | 25.0 | 26.8 | 3.6 | 60.7 | 39.3 |
| 2 | 60 | 2.93 | 30 | 61.7 | 38.3 | 28.3 | 33.3 | 23.3 | 15.0 | 40.0 | 60.0 |
| 3 | 46 | 3.03 | 29 | 52.2 | 47.8 | 37.0 | 30.4 | 21.7 | 10.9 | 65.2 | 34.8 |
| 4 | 33 | 2.91 | 20 | 48.5 | 51.5 | 39.4 | 21.2 | 24.2 | 15.2 | 63.6 | 36.4 |
| 5 | 100 | 3.14 | 37 | 60.0 | 40.0 | 30.0 | 43.0 | 23.0 | 4.0 | 58.0 | 42.0 |
| 6 | 47 | 2.93 | 27 | 72.3 | 27.7 | 36.2 | 36.2 | 10.6 | 17.0 | 57.4 | 42.6 |
| 7 | 46 | 3.14 | 23 | 60.9 | 39.1 | 21.7 | 43.5 | 28.3 | 6.5 | 58.7 | 41.3 |
| 8 | 43 | 2.84 | 24 | 41.9 | 58.1 | 39.5 | 30.2 | 23.3 | 7.0 | 74.4 | 25.6 |
| 9 | 39 | 2.98 | 26 | 46.2 | 53.8 | 23.1 | 41.0 | 20.5 | 15.4 | 51.3 | 48.7 |
| 10 | 42 | 3.04 | 22 | 71.4 | 28.6 | 19.0 | 45.2 | 28.6 | 7.1 | 71.4 | 28.6 |
| 11 | 30 | 3.12 | 17 | 73.3 | 26.7 | 13.3 | 43.3 | 16.7 | 26.7 | 36.7 | 63.3 |
| **Total** | **577** | **2.93** | **-** | **60.0** | **40.0** | **31.4** | **35.7** | **22.7** | **10.2** | **57.5** | **42.5** |

These clusters illustrate what may be represented by the neural avalanches typically shown in log-scale plots, while making explicit the spatiotemporal structure that is otherwise obscured. These clusters can be further classified into travelling (0, 2-5) and standing (1, 6-11) based on whether we observe the poles moving or not. Nevertheless, the next goal is to study to what extent these patterns can be identified in the raw, i.e., unthresholded, data. Crucially, having identified the clusters, we can subsequently ask a further question concerning their dynamics: do these patterns oscillate (i.e., appearing repeatedly with alternating polarity at their own temporal length of around 40ms), or do they remain stable, repeating with consistent polarity? This distinction maps directly onto the visible spatial structure of the clusters, as we show through the back-fitting procedure described below.

### 2.2 Back-fitting patterns to the raw data

#### 2.2.1 Validation

Due to the computational cost of using the spatiotemporal alignment method from above across the entire dataset, we opted for the correlation method (see Methods 4.6.1). To assess whether it is sufficient, we examined how well each cluster template correlates with the avalanches that originally composed it versus avalanches belonging to other clusters (Figure 2).

**Figure 2:**
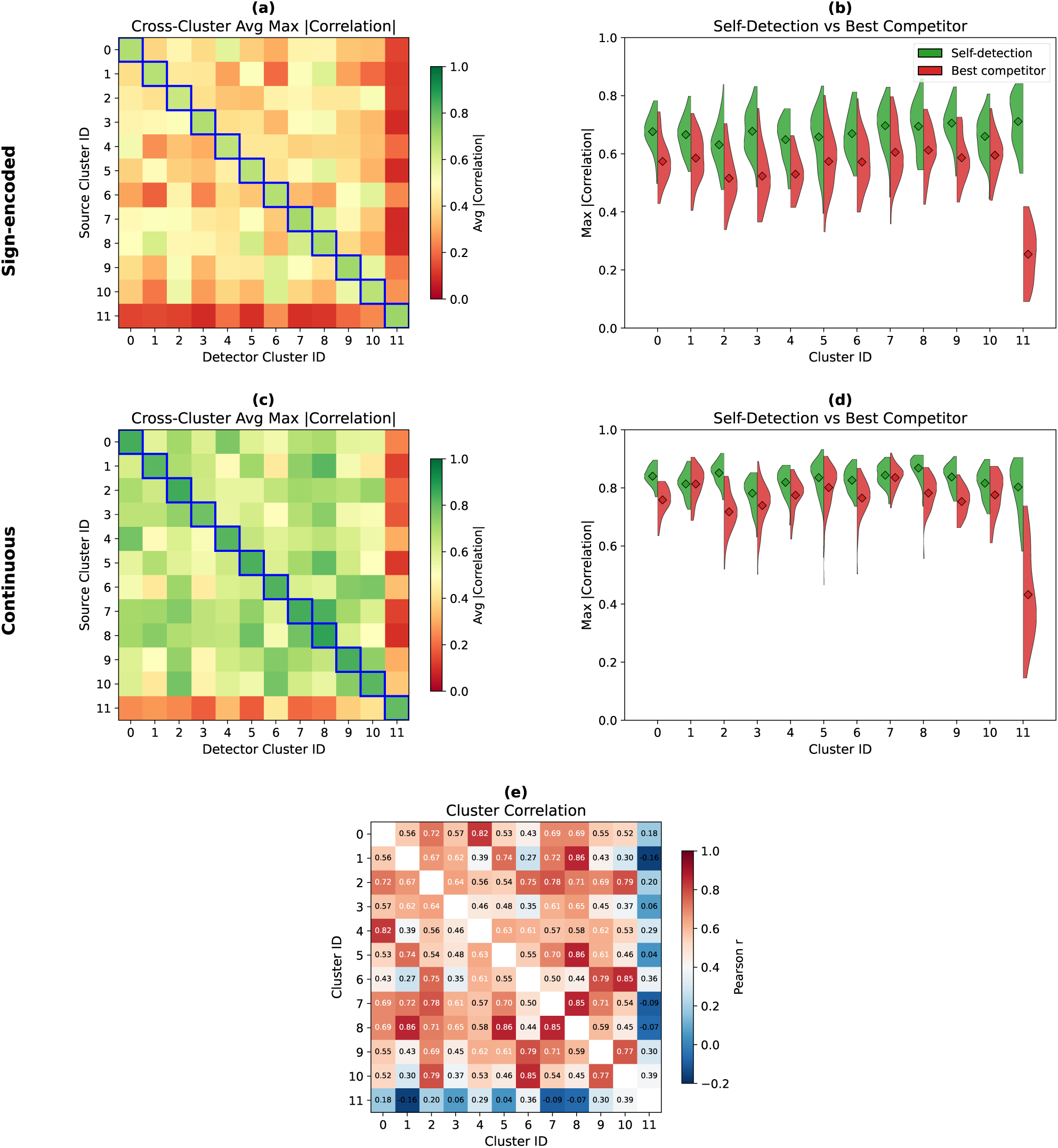
Cluster self-detection and cross-detection validation. **(a)** Cross-cluster correlation matrix for avalanche data: each cell shows the average maximum absolute correlation when using the detector cluster (column) to identify source avalanches (row). The diagonal (self-detection) is visibly higher than off-diagonal entries. **(b)** Self-detection versus best competitor for sign-encoded data: violin plots comparing, for each cluster, the maximum absolute correlation achieved by self detection (green) against the highest-scoring competing detector (red). **(c)** Cross-cluster correlation matrix for continuous avalanches, (i.e., EEG data at the same time windows as individual avalanches). The diagonal advantage is reduced, reflecting the loss of sparsity in continuous signals. **(d)** Self-detection versus best competitor for continuous data: the gap between self-detection and the best competitor narrows considerably compared to the sign-encoded case, indicating substantial overlap among cluster templates in continuous recordings. **(e)** Inter-cluster correlation matrix. Pairwise Pearson correlations between the 12 consensus cluster templates. High off-diagonal values (mean *r* = 0.532, max *r* = 0.862) reflect substantial spatial overlap among clusters distributed across the same 64-electrode array. This overlap motivated the selection of a high detection threshold (*ρ* = 0.825) to minimize spurious co-detections during backfitting.

For the sign-encoded avalanches that composed the clusters (Figure 2a,b), self-detection correlations are clearly separated from cross-cluster correlations, indicating that the correlation method can identify the correct cluster membership quite well. However, when the same time windows are evaluated using the original continuous EEG signals within the time of sign-encoded avalanches (Figure 2c,d), this separation largely disappears: cluster templates correlate highly not only with their own source avalanches but also with those from other clusters. Unlike the sign-encoded data, continuous signals contain non-zero values at every electrode and time-point, removing the sparsity that aided discrimination and making spatial overlap between cluster templates substantially more pronounced.

##### Correlation among clusters and threshold selection

This overlap raised a practical question: how distinguishable are the cluster templates from one another when used as detectors on continuous data? The inter-cluster correlation matrix (Figure 2e) reveals high pairwise correlations across most cluster pairs (mean *r* = 0.532, maximum *r* = 0.862), confirming that many clusters share considerable spatial structure despite capturing distinct propagation dynamics. To select a detection threshold *ρ* (see Pseudocode 4), we needed to balance two competing demands: the threshold must be high enough to avoid false co-detections between spatially similar clusters (Figure 2e), yet low enough to detect a sufficient number of sequences. The inter-cluster correlation matrix showed that cluster pairs with correlations above approximately one standard deviation from the mean (86th percentile, *r* ≥ 0.75) become increasingly difficult to dissociate. Averaging these high-correlation values yielded *r* = 0.811; we set the threshold slightly above this at *ρ* = 0.825. As an additional safeguard against residual overlap, we removed any detected sequences that shared more than 44 ms of temporal overlap with a sequence from a different cluster, ensuring the same neural event was not counted twice.

##### Stable and Oscillating Pattern Sequences

With the detection threshold established, we turned to the temporal dynamics of the backfitted patterns. Specifically, we asked: do successive activations of the same cluster pattern maintain consistent polarity (stable sequences) or alternate between positive and negative correlations (oscillating sequences)? As we show below, this distinction maps cleanly onto the spatial structure of the clusters themselves with the exception of cluster 8: clusters with visible spatial propagation across the scalp are precisely those that oscillate, while clusters with fixed topographic arrangements tend to be stable.

#### 2.2.2 Psilocybin is Associated with Reduced Oscillating Sequences

Psilocybin administration altered the spatiotemporal organization of avalanche sequences (Figure 3a, Table 2). After removing temporally overlapping detections—which excluded 579 sequences (16.6%), predominantly from clusters 7 and 10—we retained 2915 sequences for analysis. The overall effect of psilocybin is visible in Figure 3a. The number of oscillating sequences dropped sharply from 85.7% to 65.2%, shifting the polarity balance toward stable. Because sequences are nested within participants, we tested this shift with a within-subject permutation test rather than a pooled chi-square (see Methods); it remained significant (*p*_FDR_ = 0.022).

**Figure 3:**
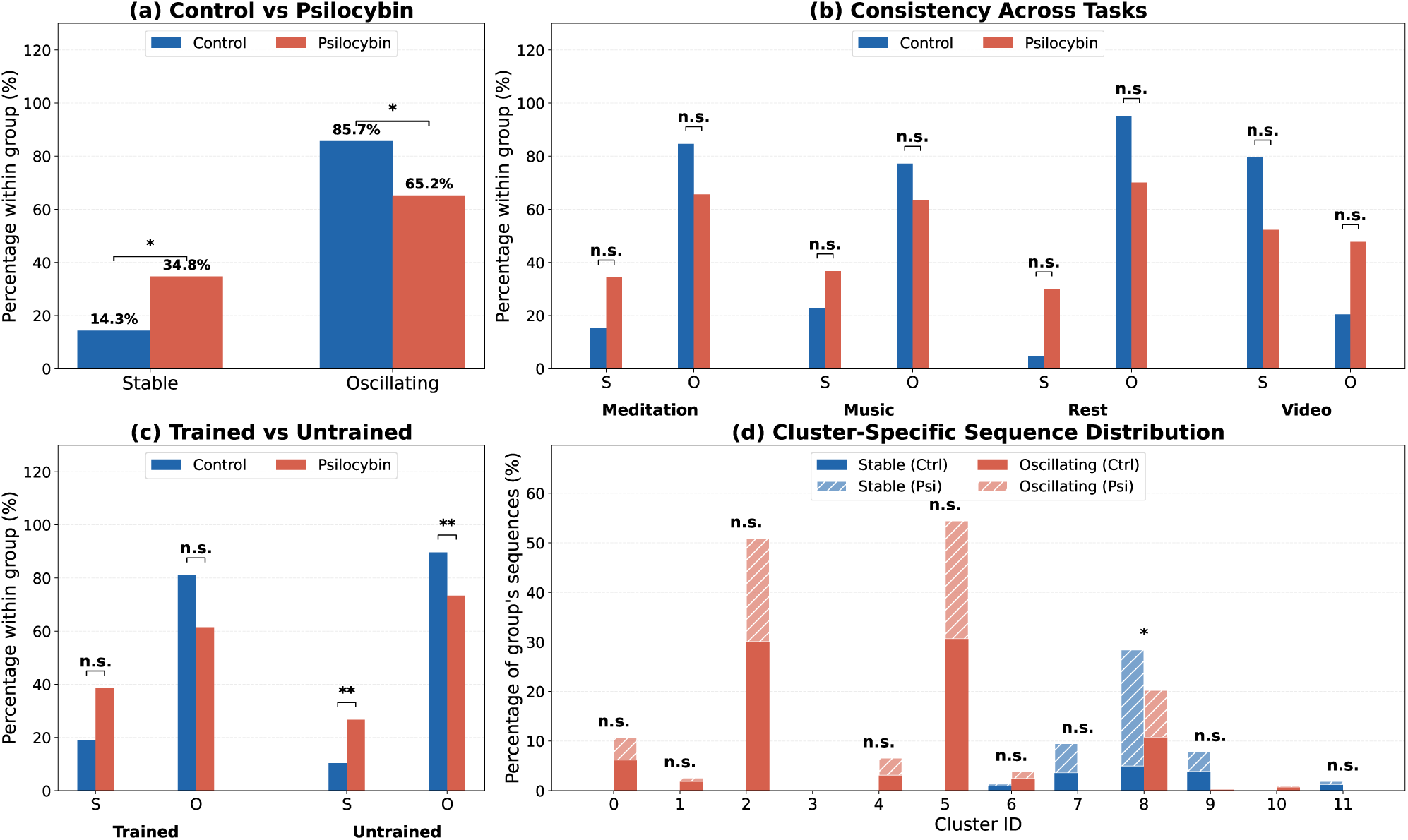
Psilocybin shifts the polarity balance of recurring sequences toward stable. Oscillating sequences are those where successive detections of the same cluster alternate in polarity while stable sequences maintain consistent polarity. Bars show within-condition proportions (each condition sums to 100%). Significance brackets show FDR-corrected subject-level permutation tests (see Methods). **(a)** Overall comparison between control and psilocybin sessions. The oscillating share (O) falls from 86% to 65% and the stable share (S) rises correspondingly; significant by within-subject permutation (*p*_FDR_ = 0.022). **(b)** Task consistency: the shift is in the same direction across tasks, but no individual task survives the permutation test (*p*_FDR_ *>* 0.1). **(c)** Drug effect within each training group: significant within untrained (*p*_FDR_ = 0.006) but not trained (*p*_FDR_ = 0.10); the groups do not differ from each other (*p*_FDR_ = 0.72). **(d)** Cluster-specific sequence counts: intrinsically oscillating clusters (notably 2 and 5, which also exhibit the clearest spatial propagation across the scalp) show the largest reductions under psilocybin, while intrinsically stable clusters (7, 9, 11, with fixed topographic arrangements) remain relatively unchanged. Cluster 8 is the only cluster exhibiting a substantial within-cluster shift from oscillating to stable dynamics under psilocybin. Significance levels are denoted as: n.s. = not significant (*p >* 0.05), ∗ *p <* 0.05, ∗ ∗ *p <* 0.01, ∗ ∗ ∗ *p <* 0.001.

**Table 2:** Cluster-specific polarity profiles reveal distinct pattern subgroups. For each cluster, the number and percentage of stable versus oscillating sequences is shown separately for control and psilocybin sessions, after removal of temporally overlapping detections (*>*44 ms). Clusters fall into three categories: intrinsically stable (7, 9, 11), intrinsically oscillating (0, 1, 2, 4, 5, 10), and mixed/drug-sensitive (6, 8). Cluster 3 contained only one sequence and is omitted. Statistical significance is assessed via within-subject permutation tests (Methods) with FDR correction; only cluster 8 showed a statistically significant within-cluster drug effect (*p*_FDR_ = 0.019).

| Cluster | Control |  |  | Psilocybin |  |  | FDR |
| --- | --- | --- | --- | --- | --- | --- | --- |
|  | Stable | Oscillating | n | Stable | Oscillating | n |  |
| *Intrinsically stable clusters* |  |  |  |  |  |  |  |
| 7 | 74 (98.7%) | 1 (1.3%) | 75 | 49 (98.0%) | 1 (2.0%) | 50 | n.s. |
| 9 | 80 (95.2%) | 4 (4.8%) | 84 | 33 (100%) | 0 (0%) | 33 | n.s. |
| 11 | 24 (100%) | 0 (0%) | 24 | 6 (100%) | 0 (0%) | 6 | — |
| *Intrinsically oscillating clusters* |  |  |  |  |  |  |  |
| 0 | 0 (0%) | 128 (100%) | 128 | 0 (0%) | 38 (100%) | 38 | — |
| 1 | 1 (2.6%) | 38 (97.4%) | 39 | 0 (0%) | 6 (100%) | 6 | n.s. |
| 2 | 1 (0.2%) | 630 (99.8%) | 631 | 0 (0%) | 171 (100%) | 171 | n.s. |
| 4 | 0 (0%) | 64 (100%) | 64 | 0 (0%) | 29 (100%) | 29 | — |
| 5 | 0 (0%) | 642 (100%) | 642 | 0 (0%) | 195 (100%) | 195 | — |
| 10 | 0 (0%) | 14 (100%) | 14 | 0 (0%) | 3 (100%) | 3 | — |
| *Mixed / drug-sensitive clusters* |  |  |  |  |  |  |  |
| 6 | 18 (26.5%) | 50 (73.5%) | 68 | 4 (25.0%) | 12 (75.0%) | 16 | n.s. |
| 8 | 103 (31.4%) | 225 (68.6%) | 328 | 192 (71.1%) | 78 (28.9%) | 270 | * |

The same directional shift appeared in meditation, music and rest tasks (Figure 3b), but no individual task survived the within-subject permutation test (all *p*_FDR_ *>* 0.1); we therefore do not interpret task-specific effects.

Video viewing ran in the opposite direction, but with only 44 sequences per condition it did not survive permutation (*p*_perm_ = 0.81) and is not interpreted.

The trained versus untrained comparison (Figure 3c) shows the shift in both groups, but it reaches significance only within untrained participants (within-subject permutation *p*_FDR_ = 0.006; trained *p*_FDR_ = 0.10), and the two groups do not differ significantly from each other (*p*_FDR_ = 0.72). The effect is thus clearest in untrained participants rather than a reliable group difference. Mindfulness-Based Cognitive Therapy (MBCT) is the training program that defines the “trained” group; see Methods.

#### 2.2.3 Cluster-Specific Polarity Dynamics

The polarity contrast was not uniform across clusters, revealing distinct subgroups of propagation patterns with intrinsic stable or oscillatory tendencies (Figure 3d; Table 2).

Clusters 7, 9, and 11 exhibited near-exclusively stable behavior in both conditions (control: 95.2–100% stable; psilocybin: 98.0–100% stable), with no statistically significant psilocybin effect on their polarity profile. These clusters maintain consistent spatial polarity across successive activations regardless of pharmacological condition; notably, their topographies show minimal spatial movement across the scalp, consistent with standing (non-propagating) patterns. Conversely, clusters 0, 1, 2, 4, 5, and 10 were near-exclusively oscillating in both conditions (97.4–100% oscillating), again with no statistically significant drug-related change in polarity. Crucially, these are precisely the clusters that show visible spatial propagation in their consensus topographies— activity moving across the scalp over the 40–44 ms avalanche window—indicating that these travelling patterns identified by clustering propagate at this temporal scale (approximately 11–12.5 Hz, alpha range; see Discussion). However, these oscillating clusters were substantially less active under psilocybin (control: 1518 sequences; psilocybin: 442 sequences), indicating that travelling wave-like patterns occur less frequently under psilocybin while their intrinsic per-cluster polarity dynamics appear unchanged.

The drug effect within a cluster appeared in cluster 8, which shifted from predominantly oscillating under control (225 oscillating vs. 103 stable, i.e. 31% stable) to predominantly stable under psilocybin (78 oscillating vs. 192 stable, i.e. 71% stable; within-subject permutation *p*_perm_ = 0.001, *p*_FDR_ = 0.019). This was the only cluster showing a statistically significant within-cluster change in polarity distribution. Cluster 8 is thus the one spatiotemporal pattern whose own dynamics change under psilocybin—becoming temporally stable (consistent polarity) rather than oscillating—and the only within-cluster effect to survive permutation, making it the most promising target for further study of how psilocybin reshapes a specific propagation mode.

Taken together, the overall shift under psilocybin arises from two complementary mechanisms: (i) lower counts of intrinsically oscillating clusters under psilocybin (primarily clusters 2 and 5, which together account for the majority of oscillating sequences), and (ii) a polarity shift within cluster 8 from predominantly oscillating to predominantly stable dynamics.

### 2.3 Summary

Backfitting cluster templates to continuous EEG recordings revealed that recurring avalanche patterns manifest as sequences of cluster activations with distinct polarity dynamics. After removing temporally overlapping detections (579 sequences, predominantly from clusters 7 and 10), psilocybin shifted the polarity balance toward stable—oscillating sequences dropped from 85.7% (1797 out of 2098) to 65.2% (533 out of 817) while stable sequences were proportionally increased under psilocybin (control: 14.3% (301); psilocybin: 34.8% (284)). This overall shift was significant at the participant level (within-subject permutation *p*_FDR_ = 0.022), whereas the per-task and trained-versus-untrained differences did not survive subject-level testing.

Individual clusters exhibited intrinsic polarity profiles: some predominantly stable (clusters 7, 9, 11), others near-exclusively oscillating (clusters 0, 1, 2, 4, 5, 10). Within-cluster polarity profiles were similar across conditions; the difference between conditions was concentrated in the count of oscillating-cluster activations. Cluster 8 was the only cluster showing a statistically significant within-cluster shift, moving from predominantly oscillating under control to predominantly stable under psilocybin (within-subject permutation *p*_FDR_ = 0.019).

## 3 Discussion

### 3.1 Summary and Spatial Symmetries of Clusters

This exploratory study presents a methodological framework for identifying and tracking recurring spatiotemporal patterns in neural avalanches from EEG recordings. The core distance computation follows the philosophy of Janati et al. (2020), who combined unbalanced optimal transport with soft-DTW for spatiotemporal alignment; we tailored their framework to EEG avalanches through the Hungarian algorithm, polarity constraints, and subsequence DTW. Applied to 64-channel EEG from 63 participants under baseline and psilocybin conditions, hierarchical clustering identified 12 distinct propagation modes. The members of these clusters did not statistically differ in being sampled from the control or psilocybin conditions which is consistent with prior microstate studies in psilocybin (Jajcay et al., 2026). Backfitting these templates to continuous recordings revealed that psilocybin shifted the polarity balance of recurring sequences toward stable, an overall shift that held within participants (subject-level permutation *p*_FDR_ = 0.022) but was not individually significant per task.

A notable property of the identified clusters is their systematic spatial symmetries. Cluster pairs 7–9, 0–4, and 2–5 show near-perfect spatial reflection across hemispheres, and within each pair, both members share the same temporal profile: clusters 7 and 9 are both stable (98.7% and 95.2% stable under control), while clusters 0 and 4 are both exclusively oscillatory. Similarly, clusters 2 and 5—which together account for the majority of detected sequences (Figure 3d)—represent mirror-identical patterns with reversed propagation direction and opposite polarity structure. Both show activity travelling across occipital-parietooccipital electrodes with 40– 44 ms periodicity. Since oscillating sequences consist of two successive detections with alternating polarity, each positive-negative pair completes one full cycle in 80–88 ms, corresponding to approximately 11–12.5 Hz. Cluster 3 starts at the left side of central electrodes and propagates in the opposite direction compared to clusters 0 and 4, suggesting not only spatial but also temporal symmetry.

These symmetries reduce the effective dimensionality of the pattern space (i.e., rather than 12 independent templates, the repertoire can be described by a smaller set of canonical patterns and their mirror transforms) and suggest that cortical dynamics operate within constrained geometric relationships. Future work should formally characterize these symmetry groups and test whether they generalize to independent datasets and recording modalities.

### 3.2 Unifying Spatiotemporal Pattern Analysis: Microstates, Travelling Waves, and Avalanches

Our findings suggest that three traditionally separate analytical frameworks—neural avalanches, EEG microstates, and travelling waves—may describe the same underlying spatiotemporal dynamics from different perspectives. The evidence for this comes from specific correspondences between our data-driven clusters and phenomena that each framework studies independently.

#### Connection to EEG microstates

EEG microstate analysis assigns every timepoint to one of 4–6 quasistable topographic states, typically explaining 60–80% of data variance (Lehmann et al., 1987; Michel and Koenig, 2018; Jajcay and Hlinka, 2023). Several of our clusters—particularly 7, 8, and 9—show spatial topographies and temporal stability remarkably similar to canonical microstate classes (Koenig et al., 2002; Britz et al., 2011; Custo et al., 2017). These are precisely the clusters classified as intrinsically stable in our analysis: they maintain fixed topographic arrangements across successive activations, with frontal-to-posterior or centroparietal organization matching the spatial structure of canonical classes. In addition, both our analysis along with microstates – although not applied in this work – allow measuring the same metrics such as occurrence, duration, coverage and transition matrix.

This correspondence is notable because it arises without any microstate-specific assumptions. Microstate analysis has been criticised for ignoring the continuous properties of EEG (Mishra et al., 2020), assuming discrete switches between states (Von Wegner et al., 2018), and forcing all data into predefined categories regardless of signal quality. Our approach addresses each of these: clusters capture time-varying spatiotemporal propagation rather than static topomaps fitted to temporal data; data segments dissimilar to any cluster are simply ignored rather than forced into a category. That microstate-like patterns emerge naturally from avalanche clustering— without assuming that the brain activity switches in discrete fashion (Michel and Koenig, 2018; Mishra et al., 2020) or requiring complete variance explanation—suggests that what microstates capture is a real feature of neural dynamics, not an artifact of the fitting procedure (Lehmann et al., 2005). At the same time, our framework goes beyond microstates by revealing what they cannot: the travelling dynamics of the remaining clusters.

#### Connection to spatiotemporal waves

Travelling waves in EEG are typically identified through dedicated detection frameworks that require prior commitments—selecting a frequency band, computing phase gradients or circular statistics, and defining coherence criteria for what constitutes directed propagation (Muller et al., 2016; Zhang et al., 2018; Bahramisharif et al., 2013; Alexander et al., 2013; Davis et al., 2021). Clusters 2 and 5, the most frequently detected patterns, emerged from purely data-driven clustering without any wave- or phase-specific assumptions, yet show clear rotating dynamics in occipital regions with alpha-frequency periodicity (2×44ms, ∼11 Hz within oscillating sequences). This methodological independence—recovering rotating wave dynamics without frequency filtering, phase estimation, or propagation criteria—further suggests that these patterns reflect genuine cortical dynamics rather than artifacts of wave-specific detection frameworks.

The alpha-frequency periodicity of these rotating patterns is consistent with travelling alpha waves documented in visual cortex (Zhang et al., 2018; Schwenk and Alamia, 2025; Mohan et al., 2024). Rotating waves have been observed in motor cortex (Rubino et al., 2006) and visual cortex (Sato et al., 2012; Benucci et al., 2007) using invasive recordings; that such patterns are detectable with the limited spatial resolution of scalp EEG suggests occipital-parietal rotating wave dynamics accessible through non-invasive methods. The systematic polarity reversals accompanying propagation direction changes add a dimension not typically captured in amplitude-based travelling wave analyses (Muller et al., 2018; Davis et al., 2021).

#### Unification through avalanches

These three frameworks—avalanches as scale-free events (Beggs and Plenz, 2003; Plenz et al., 2021), microstates as discrete topographies (Michel and Koenig, 2018), travelling waves as spatial propagation (Muller et al., 2018)—have developed largely in parallel, each with its own extraction methods, summary statistics, and theoretical motivations. Our results suggest they target overlapping aspects of the same spatiotemporal dynamics. However, by classifying avalanches based on their spatiotemporal shape, we recovered both microstate-like standing patterns (clusters 7, 8, 9) and travelling wave patterns (clusters 0, 2, 4, 5) within a single analytical framework. Backfitting then revealed that these two classes behave differently under pharmacological perturbation (under psilocybin the polarity balance shifted away from oscillating patterns and toward standing/microstate-like ones) suggesting they reflect functionally distinct modes of cortical organization.

This convergence suggests that microstate fitting and wave detection are complementary partial views of the same spatiotemporal dynamics, each foregrounding a different aspect—and that avalanche-based analysis provides a unifying framework capable of recovering both within a single pipeline.

### 3.3 Methodological Limitations

Our analysis involves multiple heuristic parameter choices: avalanche detection threshold (±3*σ*), minimum duration (11 time steps = 40 ms), spatial coherence criterion (5/8 active neighbors), clustering distance threshold (3.5), and minimum cluster size (30). These were selected through visual inspection rather than systematic optimization. Comprehensive parameter exploration is computationally prohibitive—computing the distance matrix for 10,216 avalanches required approximately 300 hours—and different choices would likely identify different patterns. However, our validation strategy partially mitigates this concern: we do not optimize parameters against an objective function but instead verify that the resulting clusters are observable in raw data through backfitting, exhibit systematic symmetries, and show consistent pharmacological modulation across independent tasks and participant groups. If heuristic choices had produced noise clusters, this convergence would be unlikely. That said, we cannot claim the methodology discovers all important spatiotemporal structures, but only that it discovers some genuine ones.

The primary technical limitation lies in backfitting. During clustering, UOT+subDTW provides flexible spatial and temporal alignment essential for grouping similar avalanches despite variation. During backfitting, correlation requires strict frame-by-frame spatial matching for computational tractability. This trade-off means the backfitting method cannot fully distinguish among cluster templates that share substantial spatial structure (Figure 2b, d). The high inter-cluster correlations (mean *r* = 0.532, max *r* = 0.862; Figure 2e) reflect genuine spatial overlap—all patterns propagate across the same 64-electrode array—but necessitated a stringent detection threshold (*ρ* = 0.825) and removal of temporally overlapping detections. Improving backfitting through tractable approximations to UOT+subDTW for continuous data remains an important open problem. Nevertheless, the current approach is sufficient to reveal systematic pharmacological modulation, suggesting the detected patterns carry meaningful signal despite imperfect discrimination.

A natural question is whether surrogate analyses are needed to validate the clusters. We argue the clusters are already validated by their internal consistency: bilateral mirror symmetries, clean separation into stable and oscillating profiles, and consistent drug effects across tasks. Purely random surrogates would not reproduce this structure. The more informative test would be surrogates preserving both spatial and temporal correlations simultaneously—essentially a linear spatiotemporal model of the EEG (Jajcay and Hlinka, 2023). If the same clusters emerged from such surrogates, it would indicate that our nonlinear pipeline captures structure already explained by linear correlations. This would not invalidate the clusters but would question whether UOT+subDTW is necessary. We leave this comparison for future work. Finally, the 64-electrode montage provides limited spatial resolution, and volume conduction may contribute to apparent spatiotemporal structure; higher-density EEG or source-reconstructed data would help address this.

### 3.4 Conclusion

We introduced a framework for comparing and clustering individual neural avalanches by their spatiotemporal shape, and applied it to 64-channel EEG from 63 participants under psilocybin and baseline conditions. Hierarchical clustering revealed 12 recurring propagation modes with systematic bilateral symmetries—mirror-symmetric cluster pairs sharing the same polarity dynamics—suggesting that cortical dynamics operate within constrained geometric relationships. Backfitting these templates to continuous recordings showed that psilocybin shifted the polarity balance of recurring sequences toward stable patterns and away from oscillating ones. The biggest driver of this effect is specifically localised to cluster 8, the single pattern whose own polarity is significantly reorganised by psilocybin (31% to 71% stable, the only per-cluster effect surviving permutation) and the most promising target for future study, as we can specifically see which neural dynamics changes the temporal structure.

The approach recovers both microstate-like standing patterns and travelling wave dynamics within a single data-driven pipeline—without assuming discrete state switches, frequency bands, or propagation criteria. This suggests that microstate analysis and travelling wave detection are not independent windows onto different phenomena, but complementary partial views of the same spatiotemporal dynamics; avalanche-based analysis provides a more general lens capable of capturing both simultaneously.

## 4 Methods

### 4.1 Dataset and Participants

#### 4.1.1 PsiConnect Study Design

Data for this study were drawn from the PsiConnect study (Novelli et al., 2026). The dataset comprises EEG recordings from 63 participants (after preprocessing) under psilocybin and baseline administration, measured in four different tasks (resting state, video viewing, music listening, and meditation), divided into two groups based on meditation training. The mindfulness-trained group (n=33; mean age 35.52 years, SD=10.64, range 20–54; 14 female, 1 non-binary) completed an eight-week Mindfulness-Based Cognitive Therapy (MBCT) program (Williams et al., 2011) involving daily meditation practice (approximately 20 minutes) and weekly 70-minute group sessions. The untrained group (n=30; mean age 40.03 years, SD=10.85, range 19–53; 16 female) received no meditation training. Group assignment was non-randomised but balanced for demographic characteristics.

#### 4.1.2 Experimental Protocol

Each participant underwent two EEG recording sessions: a baseline session conducted approximately one week prior to psilocybin administration, and an acute session performed 75–120 minutes following ingestion of 19 mg psilocybin under medical supervision. Within each EEG session, participants completed four sequential tasks in the following order: naturalistic movie viewing (5 minutes), eyes-closed resting state (5 minutes), guided meditation (5 minutes), and music listening (actual duration varied between 320–340 seconds depending on the recording).

#### 4.1.3 Data Acquisition

Continuous EEG was recorded using a 64-channel BrainAmp MR Plus amplifier system following the international standard 10-20 electrode placement (HH, 1958), which positions electrodes at intervals of 10% and 20% of skull dimensions. Data were acquired at 500 Hz sampling rate with the frontal midline electrode (FCz) serving as the online reference. Participants remained seated in an upright, comfortable position throughout recordings. All procedures received institutional ethical approval and participants provided written informed consent.

#### 4.1.4 Preprocessing Pipeline

Raw EEG data underwent a multi-stage preprocessing workflow implemented in EEGLAB (Delorme and Makeig, 2004):

##### Initial filtering and referencing

Continuous data were re-referenced to the average of all electrodes to facilitate topographic analysis (Murray et al., 2008; Dien, 1998) and band-pass filtered between 1–40 Hz to isolate neural oscillations while removing slow drifts and high-frequency noise.

##### Automated artifact rejection

The CleanRawData plugin identified and removed noisy channels and time segments based on statistical thresholds, followed by Artifact Subspace Reconstruction (ASR) to correct remaining transient artifacts through subspace projection methods. Recordings with excessive data loss (*>*25% channels removed or *>*66% time segments rejected) were excluded from analysis. The removed channels are later on interpolated for proper identification of spatiotemporal structures.

##### Independent component analysis

ICA decomposition isolated spatially fixed sources of activity, enabling automated identification and removal of stereotyped artifacts (ocular movements, muscle activity, electrode noise) using the ICLabel classifier (Pion-Tonachini et al., 2019).

##### Quality verification

All preprocessed datasets underwent visual inspection using MICROSTATELAB’s k-means clustering diagnostic tool (Nagabhushan Kalburgi et al., 2023) to find whether the noisy electrodes were being clustered. If so, the ICA was checked again.

##### Final output

For computational efficiency in subsequent avalanche detection, retained datasets were down-sampled to 250 Hz, preserving temporal dynamics while reducing computational demands. Table 3 summarises the final dataset composition across experimental conditions, including the mean recording duration for each condition. Recording durations were generally consistent across conditions (4-5 minutes), with video-viewing sessions showing slightly shorter durations (mean: 3.8-4.1 minutes) compared to other tasks (mean: 4.4-5.3 minutes). Music is naturally longer than the rest due to the original recording of 7 minutes, compared to 5 minutes in the other tasks. Data loss occurred predominantly in the video-viewing condition during psilocybin sessions, likely reflecting increased eye motion artifacts during acute psychedelic effects.

**Table 3:** Dataset availability and recording duration. Mean ± standard deviation of recording duration (in minutes) and number of datasets (n) retained after preprocessing for each group (Control: no meditation training; MBCT: mindfulness-based cognitive therapy participants), experimental session (Baseline vs. Psilocybin), and task condition.

| Session | Group | Video | Rest | Meditation | Music |
| --- | --- | --- | --- | --- | --- |
| Baseline | Untrained | 4.0±0.8 (n=24) | 4.9±0.1 (n=28) | 4.8±0.4 (n=29) | 5.3±0.3 (n=29) |
|  | Trained | 4.1±0.7 (n=29) | 4.8±0.5 (n=31) | 4.7±0.6 (n=31) | 5.0±0.7 (n=31) |
| Psilocybin | Untrained | 3.9±0.7 (n=17) | 4.6±0.5 (n=25) | 4.6±0.4 (n=26) | 4.9±0.7 (n=24) |
|  | Trained | 3.8±0.7 (n=22) | 4.4±0.6 (n=27) | 4.5±0.6 (n=28) | 4.9±0.7 (n=28) |

This preprocessing pipeline ensured that subsequent avalanche analyses operated on relatively clean neural signals while preserving the spatiotemporal structure necessary for identifying propagation patterns.

### 4.2 Avalanche Detection and Preprocessing

#### 4.2.1 Avalanche Identification

Neural avalanches were identified using a threshold-based approach following established methods in the field (Beggs and Plenz, 2003; Plenz et al., 2021). For each electrode per each dataset, the signal was z-score normalised (mean-centered and divided by standard deviation), shown in Figure 4(a). Time points were marked as active when at least one electrode exceeded a threshold of ±3 standard deviations, and the avalanche ended when no electrode remained active above the threshold. Continuous periods of supra-threshold activity were identified as candidate avalanches, with a minimum duration requirement of 11 time steps (40 ms at 250 Hz) to focus on sustained, informative avalanche events. For each avalanche, only electrode-time points exceeding the ±3 standard deviation threshold were retained with sub-threshold values set to zero. This is visualised in Figure 4(b).

**Figure 4:**
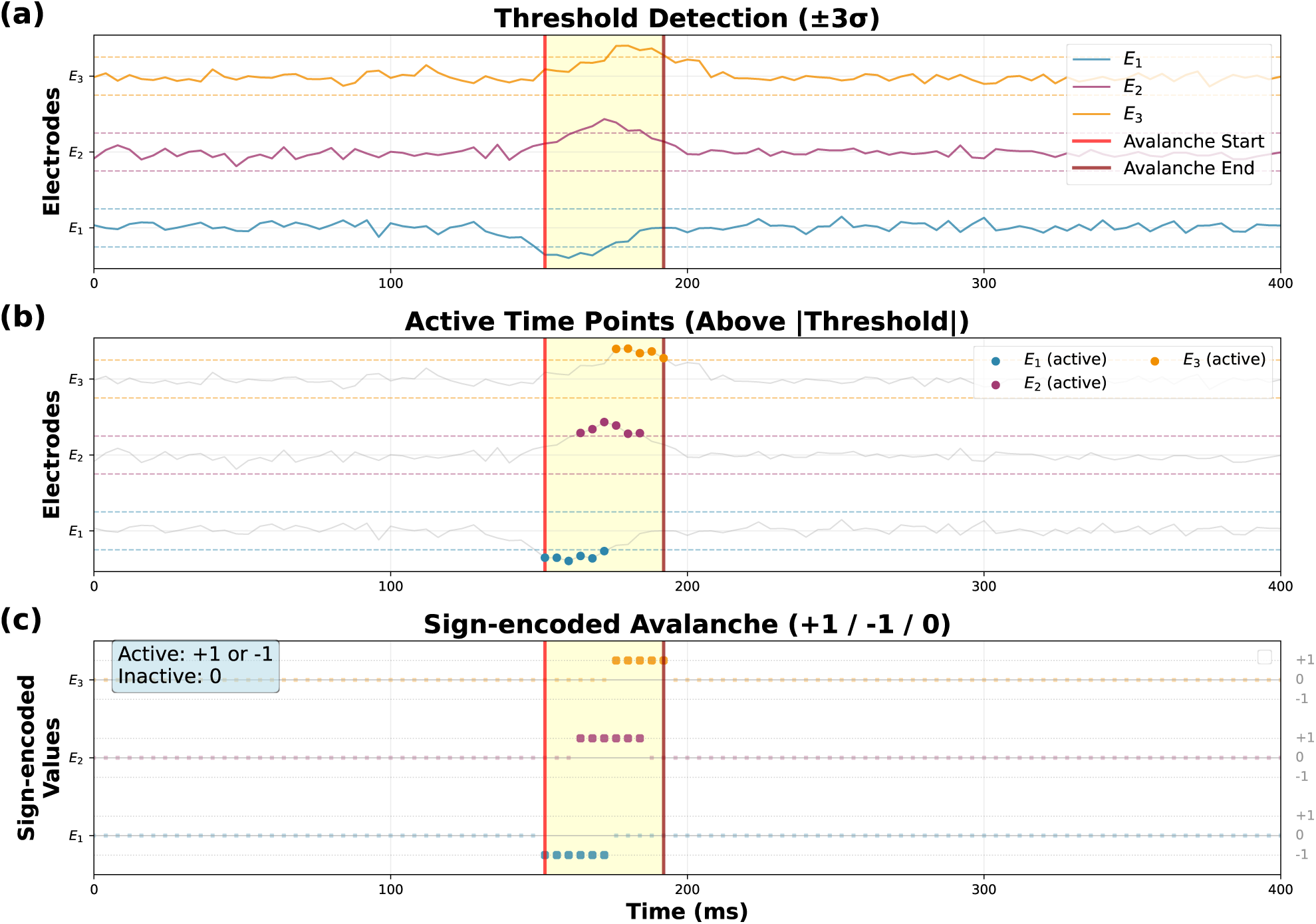
Neural avalanche identification process. The figure illustrates the three-step procedure for detecting and characterizing neural avalanches from multi-electrode recordings. **(a) Threshold Detection:** Raw signals from three electrodes (*E*_1_, *E*_2_, *E*_3_) are z-score normalised, with each electrode’s trace vertically offset for clarity. Dashed horizontal lines indicate the detection thresholds at ±3*σ* for each electrode. The yellow-shaded region marks the detected avalanche period, bounded by red vertical lines indicating avalanche start and end times. During the avalanche, *E*_1_ (blue) crosses the negative threshold while *E*_2_ (purple) and *E*_3_ (orange) cross the positive threshold. **(b) Active Time Points:** Time points where electrode signals exceed the absolute threshold (|threshold| = 3*σ*) are highlighted as colored dots, shown against the full signal traces (gray). The yellow-shaded region and red vertical lines mark the avalanche boundaries. Only supra-threshold activity is retained for avalanche analysis. **(c) Sign-encoded Avalanche:** Detected avalanche activity is sign-encoded while preserving polarity: positive threshold crossings are set to +1, negative crossings to -1, and sub-threshold activity to 0. *E*_1_ exhibits negative polarity (blue dots below zero line) while *E*_2_ and *E*_3_ show positive polarity (dots above their respective zero lines). The yellow-shaded region and red vertical lines continue to mark the avalanche boundaries across all panels.

#### 4.2.2 Spatial Filtering for Propagating Avalanches

By visually checking avalanches, some of them showed artifacts, such as a single electrode active for several timesteps, or robust activity around jaws. Thus, to focus our analysis on avalanches exhibiting spatially coherent propagation rather than random fluctuations, we applied a spatial filter. For each avalanche, we examined every time point and identified electrodes with active (non-zero) values. For removing noisy avalanches (e.g., the ones where a single electrode is active across several timepoints) an avalanche was retained if, at any time point, at least one active electrode had five or more of its eight nearest spatial neighbors (i.e., electrodes) also active with the same polarity. This had to be fulfilled twice for both polarities, and the electrodes creating the edge of the EEG device were ignored, apart from occipital and parietooccipital electrodes. This heuristic filtering assures that avalanches of pure artifacts would not be considered for analysis, while preserving also those avalanches that show at least some burst of activity.

#### 4.2.3 Peak Representation

Following spatial filtering, avalanche amplitudes were converted to a sign-based representation. Positive amplitude values were mapped to +1, negative values to -1, and zeros remained 0. This sign-encoded representation preserves the spatial and temporal structure of activation patterns while normalizing amplitude differences across electrodes and participants, making avalanches directly comparable for pattern-based analysis. This encoding is the final step shown in Figure 4(c).

### 4.3 Spatiotemporal Alignment and Distance Computation

Now, with avalanches identified, this section focuses on finding distances from avalanche to avalanche, in order to properly cluster the similar ones together.

#### 4.3.1 Spatial Distance Between Activation Patterns

For each pair of time points from two avalanches, we computed the spatial distance between their activation patterns using unbalanced optimal transport (UOT) with polarity constraints. Electrodes with positive activation (+1) could only be matched to other positive activations, and negative activations (-1) only to other negatives. Within each polarity group, we solved the optimal assignment problem using the Hungarian algorithm with electrode positions – with their x, y and z coordinates on the head – as the ground metric for distance. Unmatched electrodes (due to cardinality imbalance) were penalised by their minimum possible distance to any electrode in the opposing pattern. This minimises the punishment of overlapping activities that might have been thresholded differently. For a more intuitive understanding of this process, see Figure 5. Finally, we normalize the distance per electrode, limiting the penalization of having more active electrodes, which ultimately leads to higher transport. The entire procedure is described in Pseudocode 1. Also, note that for each avalanche, we reversed the polarities, doubling the number of avalanches we worked with. This follows the logic to provide negative correlations as well as positive ones for the back-fitting, that will be described later on.

**Figure 5:**
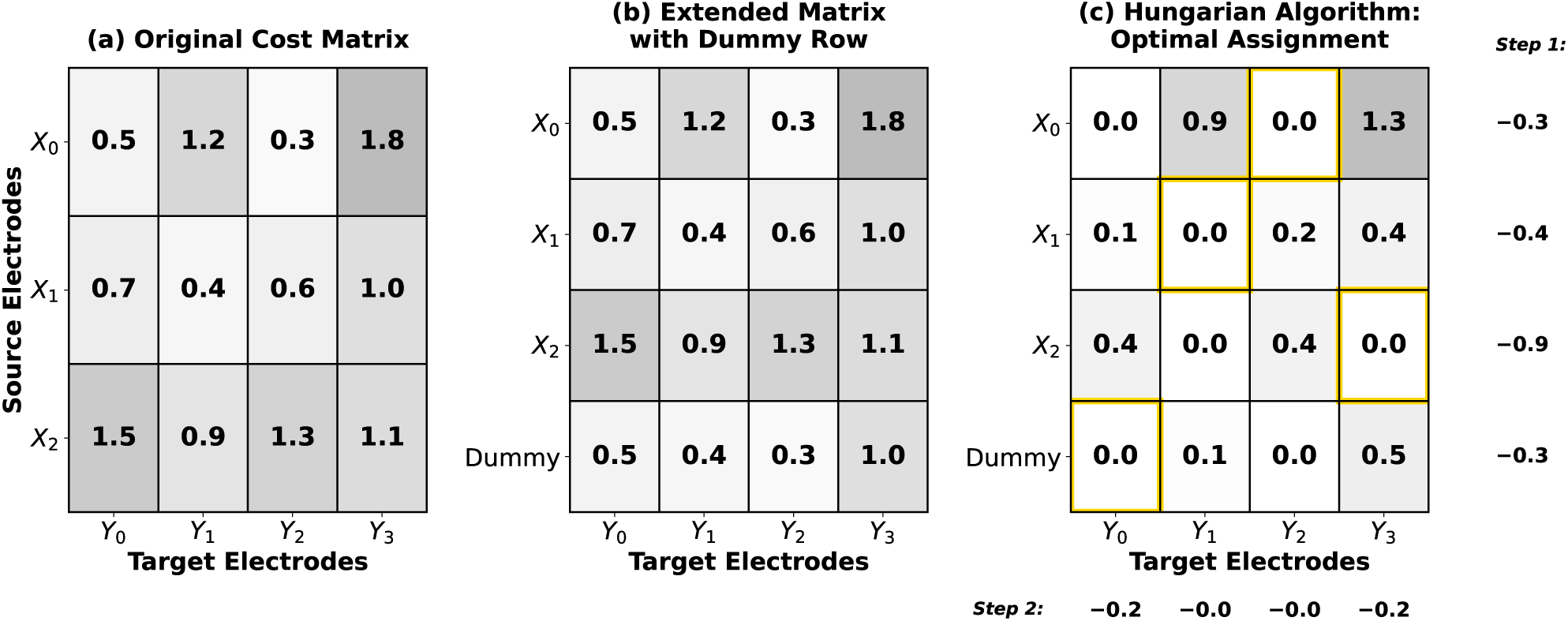
Hungarian Algorithm for Spatial Pattern Alignment with Cardinality Imbalance. The linear sum assignment problem is solved through matrix reduction to find optimal electrode matching between spatial patterns. **(a) Original Cost Matrix:** *X* and *Y* represent spatial patterns with positive or negative activity values from the dataset. The matrix entries contain normalised Euclidean distances between electrode positions, where each element *d_ij_* represents the physical distance from source electrode *X_i_* to target electrode *Y_j_*. In this example, pattern *X* has 3 active electrodes while pattern *Y* has 4, creating a cardinality imbalance. **(b) Extended Matrix with Dummy Row:** To handle the imbalance, the matrix is extended by adding a dummy row. Each dummy row entry is set to the minimum value of its corresponding column: dummy*_j_* = min*_i_*(*d_ij_*), allowing unmatched target electrodes to be assigned to the dummy with minimal penalty. **(c) Hungarian Algorithm and Optimal Assignment:** The algorithm proceeds in two steps: *Step 1* subtracts the minimum value from each row (min*_j_*(*d_ij_*) for row *i*), then *Step 2* subtracts the minimum value from each column 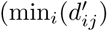 for column *j*). Cells containing zeros after reduction indicate potential optimal assignments (highlighted in gold). The final optimal assignment minimises total transport cost, calculated as the sum of original matrix values at assigned positions: 0.3 + 0.4 + 1.1 + 0.5 = 2.3. This approach ensures both polarity-matched assignments and handles cardinality imbalances through dummy assignments.

#### 4.3.2 Temporal Alignment via Subsequence DTW

Once the UOT has been computed, we followed with subsequence Dynamic Time Warping (subDTW), which finds the best temporal alignment (i.e., the lowest distance path), resulting in a single scalar value representing the final distance between two avalanches. subDTW particularly allows that the shorter avalanche aligns to any contiguous subsequence of the longer avalanche, with adaptive penalties (0.1% of minimum non-zero cost) on insertions and (2*0.1% of minimum non-zero cost) on deletions to discourage excessive warping while allowing necessary temporal flexibility. These penalties favor diagonal moves in the alignment path, preferring paths of length at least *N* (the length of the shorter sequence) to maintain one-to-one temporal correspondence. If the optimal alignment *i^∗^* yields a path shorter than N, we select the next-best alignment (second-minimum in *D*[*i^∗^, N*]) and continue until the length requirement is satisfied. This procedure is described in Pseudocode 2, and the visualization of this procedure with the matrix from the Pseudocode 1 in Figure 6.

**Figure 6:**
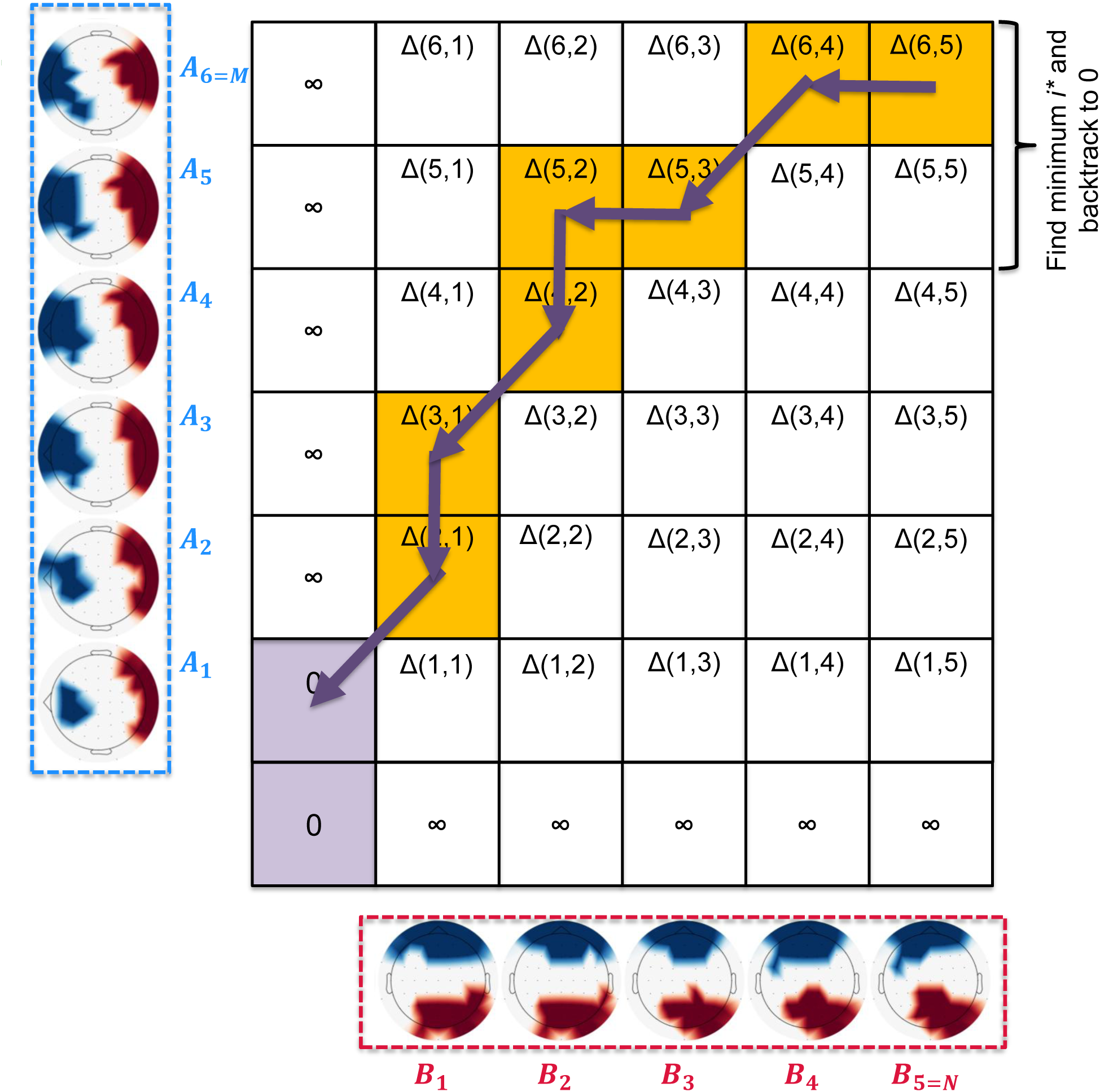
This figure demonstrates how subDTW finds the optimal alignment between avalanche sequences of different durations, allowing the shorter sequence to match a contiguous portion of the longer sequence while maintaining temporal correspondence and minimizing the total alignment cost.

##### Algorithm 1

Spatial Alignment via UOT

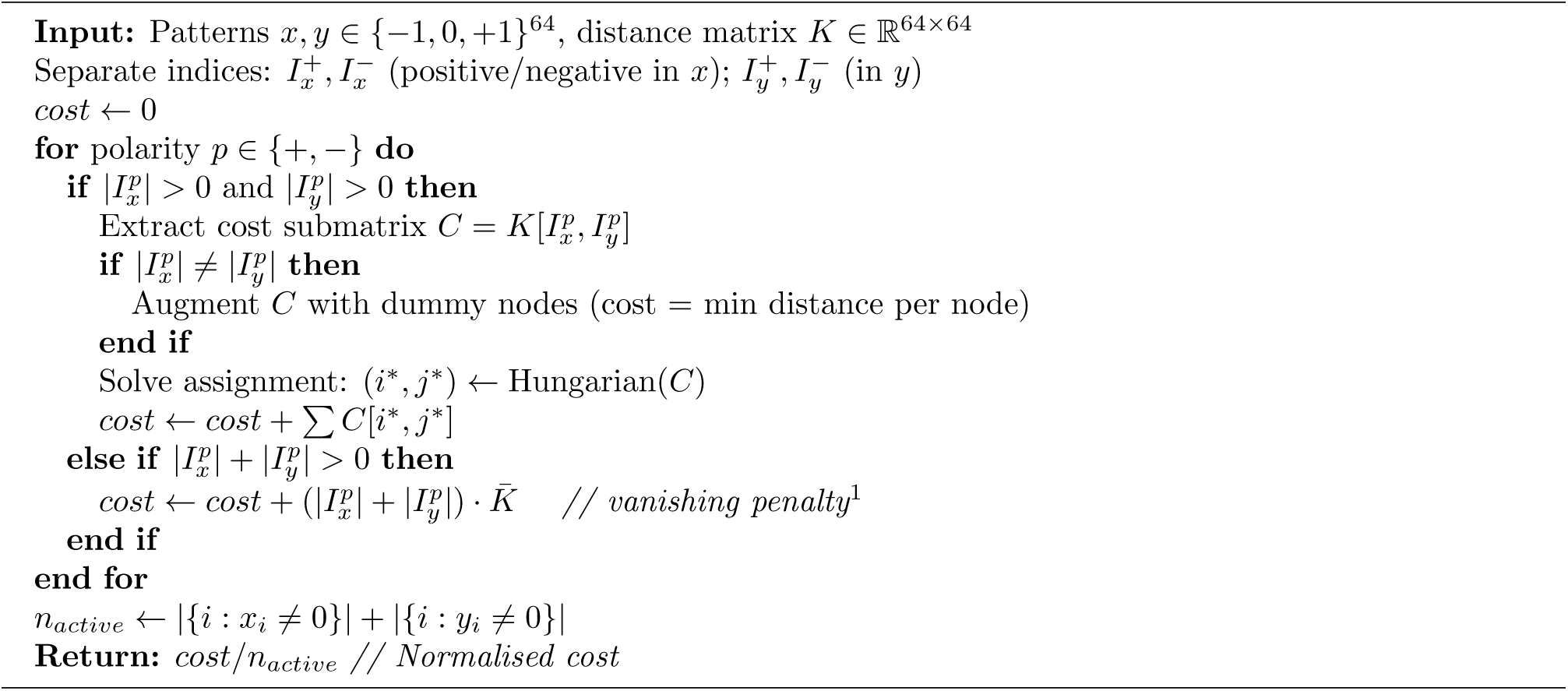

##### Algorithm 2

Subsequence DTW Alignment

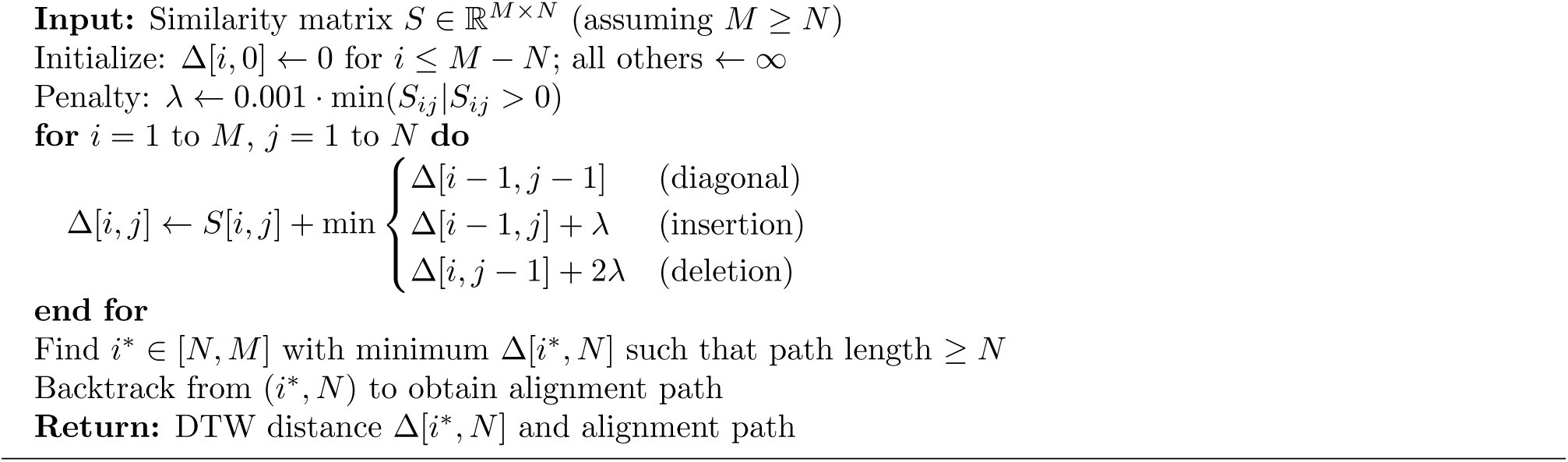

#### 4.3.3 Summary

This two-stage approach—optimal transport for spatial alignment, followed by subDTW for temporal alignment— yields a distance metric between avalanches that respects both their spatial structure and temporal dynamics while accommodating differences in duration and timing.

This method provides a distance matrix from avalanche to avalanche, where the single subDTW scalar value represents the distance of two avalanches. Achieving this, the next step is to cluster them.

### 4.4 Hierarchical Clustering

To identify recurring avalanche patterns across participants, tasks, and drug conditions, we applied hierarchical clustering. The algorithm operates as follows: We compute the average-linkage hierarchical clustering on the pairwise distance matrix of all avalanches. The resulting dendrogram is cut at a fixed distance threshold (*t* = 3.5), producing an initial set of clusters. We then filter these clusters by size, retaining only those containing at least *n*_min_ = 30 avalanches. Avalanches in clusters below this threshold are labeled as noise. Each retained cluster is characterised by its size (number of avalanches) and tightness (average internal pairwise distance).

The threshold was chosen by visual inspection of the dendrogram; no automated criterion was used. The choice of 30 avalanches within a cluster was chosen to work with only the most frequently appearing avalanches, that also show a strong average avalanche based on the medoid identification, as explained next.

### 4.5 Medoid Identification and Consensus Alignment

For each identified cluster, we constructed a representative consensus spatiotemporal pattern by aligning all member avalanches to a common temporal reference. Rather than arbitrarily selecting a reference or attempting to construct an average timeline from scratch, we used the cluster medoid – the avalanche with minimum summed distance to all other cluster members – as the natural reference timeline. This medoid-based approach ensures that the consensus pattern is grounded in an actual observed avalanche rather than a potentially artificial aggregate. Due to the subDTW algorithm and the artificial choice of 11 time steps (i.e., 40ms), most of the medoids have the same temporal size.

The consensus construction proceeds in three stages: medoid identification, warping to medoid timeline, and averaging. First, for each cluster, we compute the medoid by summing the pairwise distances from each avalanche to all others within the cluster and selecting the avalanche with the minimum sum. This medoid serves as the temporal scaffold for the consensus pattern. Second, we warp each avalanche in the cluster to align with the medoid’s timeline using the precomputed subDTW paths. For each time point in the medoid, we identify all corresponding frames in the target avalanche according to the alignment path. When multiple frames map to a single medoid time point (temporal compression), we average them. When medoid frames have no corresponding frames in the target avalanche (due to alignment endpoints), they receive zero contribution from that avalanche. Third, we compute the consensus pattern by averaging the warped avalanches across all cluster members, weighted by the number of avalanches contributing at each time point. This is written below in Pseudocode 3:

#### Algorithm 3

Medoid-Based Consensus Alignment

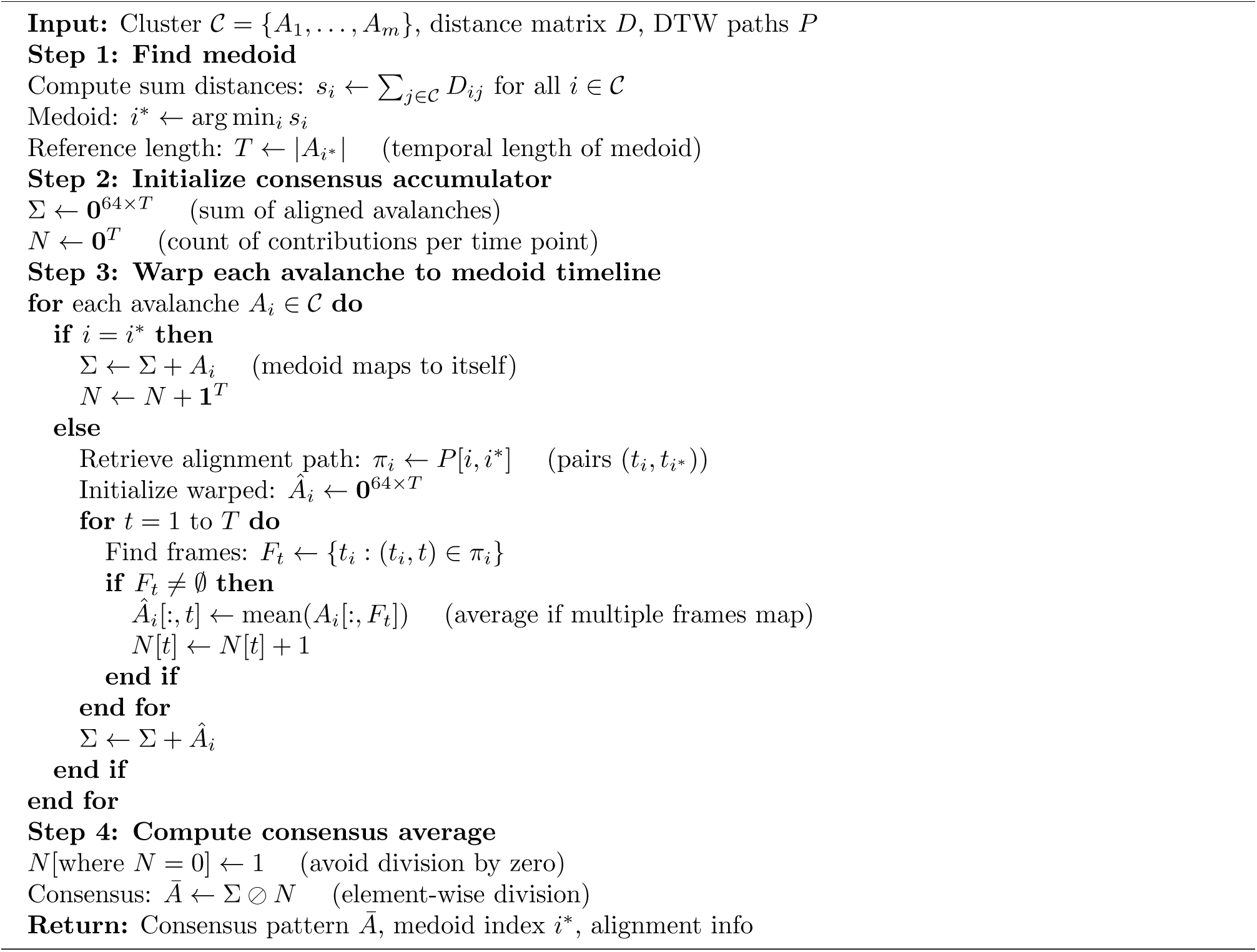

### 4.6 Spatiotemporal Correlation for Pattern Matching

The goal of this section is to find these cluster avalanches in the original dataset by back-fitting. To quantify the similarity between cluster templates and ongoing neural activity, we used a spatiotemporal correlation method. Our approach aimed to capture spatially distributed patterns that may exhibit slight variations in electrode-level activity while maintaining overall structure.

#### 4.6.1 Frame-by-Frame Spatiotemporal Correlation

We computed the spatiotemporal correlation between a cluster template pattern **P** ∈ ℝ*^n×T^* and a data window **D** ∈ ℝ*^n×T^*, where *n* is the number of electrodes and *T* is the number of timepoints. The key innovation is to compute correlation frame-by-frame. This captures both the spatial structure at each moment and how that structure evolves over time. This is described in Pseudocode 4 below:

##### Algorithm 4

Vectorised Spatiotemporal Correlation

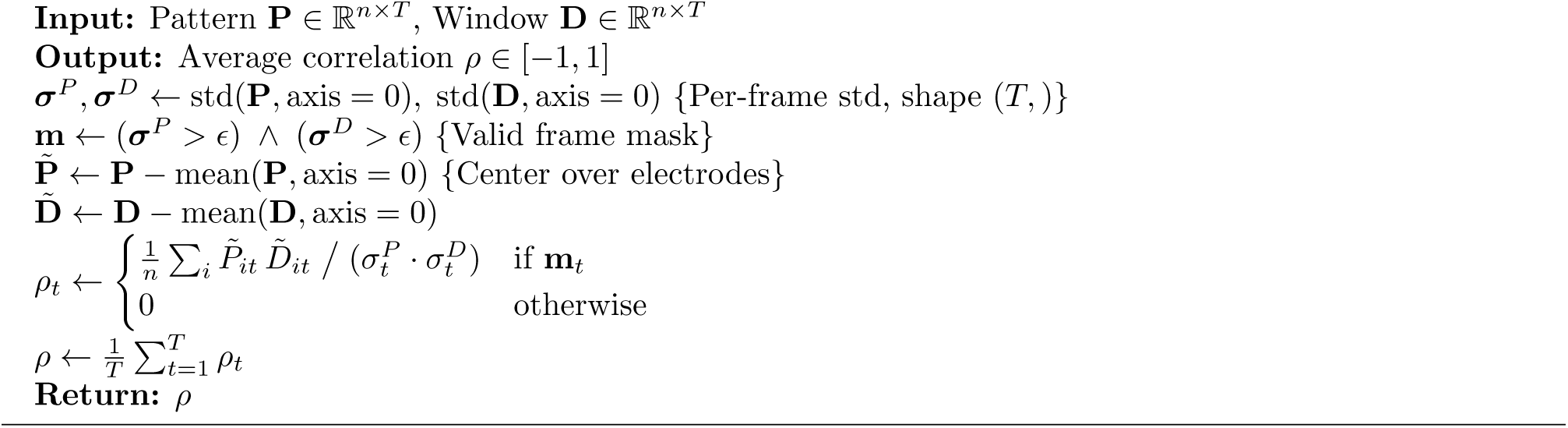

Finally, we apply this spatiotemporal correlation method for each dataset and each cluster pattern. We then slid throughout the data by each timestep where we computed the correlation at each position.

#### 4.6.2 Integration with Sequence Detection

The spatiotemporal correlation method produces continuous timeseries of correlation values for each cluster pattern. These correlation values then serve as input to the sequence detection algorithm (described in the next section), which identifies sequences of high-correlation activations that occur in rapid succession. By first computing spatiotemporal correlations and then detecting sequences in these correlation timeseries, we can characterize both the presence of recurring patterns and their temporal clustering properties. This framework allows us to quantify how recurring avalanche patterns manifest in continuous neural recordings and how their temporal dynamics are modulated by experimental manipulations such as psilocybin administration.

### 4.7 Sequence Detection and Temporal Dynamics

To characterize rapid repetitive activation patterns, we implemented a sequence detection algorithm that identified sequences of cluster activations occurring in close temporal proximity. A sequence is defined by at least two cluster activations (configurable, default: 2) with correlation of at least |*ρ*| = 0.825 with a tolerance of 1 frame delayed.

### 4.8 Statistical Analysis

Detected sequences are nested within participants: a single participant contributes many sequences, and those sequences are not independent. Pooling them into one 2 × 2 contingency table (polarity × condition) and applying a standard chi-square test therefore commits pseudoreplication—it treats dependent observations as independent, inflating the effective sample size from the number of participants to the number of sequences and yielding strongly anti-conservative *p*-values. We retain the pooled chi-square *value* as a descriptive statistic but assess significance with permutation tests that exchange labels at the participant level, so that the null distribution respects the design.

For the drug contrast (overall, per task, and per cluster), control versus psilocybin is a within-subject crossover factor. We therefore build the null by independently flipping each participant’s control/psilocybin label with probability 0.5 and recomputing the pooled chi-square; only participants present in both arms of a given slice contribute, and single-arm participants are dropped. For the between-subject trained-versus-untrained contrast, the null is built by shuffling the training label across participants while each participant keeps its own sequence counts. In both cases we draw 10,000 permutations and compute a two-sided *p*-value as 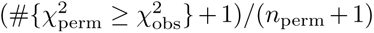, with the added unit ensuring a strictly positive, exact Monte-Carlo *p*-value. Families of tests are corrected with the Benjamini–Hochberg FDR procedure.

Finally, because the two conditions differ markedly in the total number of detected sequences (control: 2098; psilocybin: 817), absolute counts are not directly comparable across conditions. All polarity comparisons and the corresponding figure panels are therefore expressed as within-condition proportions (each condition normalised to sum to 100%), which isolates the change in polarity *balance* from the difference in overall sequence yield.

## Acknowledgments

This study is a result of the research funded by the was supported by the Czech Ministry of Education, Youth and Sports under the OP JAK grant CZ.02.01.01/00/23 025/0008715 and by the Czech Science Foundation projects No. 21-32608S and No. 23-07074S.

## Data Availability

The full analysis package is available as a stppy package at: https://github.com/filipnovicky/stppy/. The PsiConnect study with all datasets is available here: https://openneuro.org/datasets/ds006110/versions/1.2.0.

## Technical Terms

**Neural Avalanche** A burst of brain activity that begins when signals exceed a statistical threshold and ends when activity returns to baseline.

**Optimal Transport** A mathematical framework for finding the minimum-cost way to rearrange one spatial distribution of mass into another.

**Hungarian Algorithm** An efficient method for solving assignment problems, here used to optimally match active electrodes between two brain activity patterns.

**Dynamic Time Warping** A technique that aligns two time series of different lengths by stretching or compressing time to minimize their overall distance.

**Subsequence DTW** A variant of dynamic time warping that finds the best-matching portion within a longer sequence for a shorter query sequence.

**Hierarchical Clustering** An unsupervised method that groups data by iteratively merging the most similar pairs, producing a tree-like structure of nested clusters.

**Medoid** The most centrally located member of a cluster, minimizing total distance to all other members; used as a representative example.

**Travelling Wave** A spatiotemporal pattern in which a peak of neural activity propagates systematically across the cortical surface over time.

**EEG Microstates** Brief periods (60–120 ms) during which the scalp voltage topography remains quasi-stable before transitioning to a different configuration.

## Footnotes

1 If no electrodes of a given polarity exist in either pattern, all such electrodes are considered having a cost of the mean pairwise electrode distance.

